# Genetic variants affect diurnal glucose levels throughout the day

**DOI:** 10.1101/2024.07.22.604631

**Authors:** Nasa Sinnott-Armstrong, Satu Strausz, Lea Urpa, Erik Abner, Jesse Valliere, Estonian Biobank Research Team, Priit Palta, Hassan S Dashti, Mark Daly, Jonathan K Pritchard, Richa Saxena, Samuel E Jones, Hanna M Ollila

**Author notes:** Equal contribution (Samuel E Jones, Hanna M Ollila). Estonian Genome Center, Institute of Genomics, University of Tartu, Tartu, Estonia. List of Estonian Biobank Research Team is available as part of the supplementary information., Estonian Biobank.

## Abstract

Circadian rhythms not only coordinate the timing of wake and sleep but also regulate homeostasis within the body, including glucose metabolism. However, the genetic variants that contribute to temporal control of glucose levels have not been previously examined. Using data from 420,000 individuals from the UK Biobank and replicating our findings in 100,000 individuals from the Estonian Biobank, we show that diurnal serum glucose is under genetic control. We discover a robust temporal association of glucose levels at the Melatonin receptor 1B (*MTNR1B)* (rs10830963, P = 1e-22) and a canonical circadian pacemaker gene Cryptochrome 2 (*CRY2)* loci (rs12419690, P = 1e-16). Furthermore, we show that sleep modulates serum glucose levels and the genetic variants have a separate mechanism of diurnal control. Finally, we show that these variants independently modulate risk of type 2 diabetes. Our findings, together with earlier genetic and epidemiological evidence, show a clear connection between sleep and metabolism and highlight variation at *MTNR1B* and *CRY2* as temporal regulators for glucose levels.

## Introduction

Circadian rhythms not only orchestrate the timing of wake and sleep but also regulate homeostasis within the body. In addition, environmental cues, such as light, temperature, feeding, and exercise modulate circadian rhythms. Circadian rhythms are controlled by core circadian genes *CLOCK* and *BMAL1* and their repressors *PER* and *CRY* genes^1^. This loop is synchronized daily by light, temperature, feeding and activity. Circadian rhythms also significantly contribute to physiological processes, including diurnal variation in homeostasis and gene expression^2, 3^.

Melatonin, a molecule secreted at night from the pineal gland, is an important factor in regulating diurnal rhythms in humans. Its role extends from regulating circadian rhythms^4^ and circadian gene expression to influencing the timing and structure of sleep^5, 6^. Moreover, melatonin is routinely measured in clinical settings for diagnosis of suspected sleep-wake disorders. Additionally, it is commonly administered orally to adapt to time zones due to its phase-shifting properties and to treat insomnia due to its mild sedative effects.

Moreover, sleep and metabolism are tightly linked through diurnal rhythms^7^. Disrupted metabolism, exemplified by high glucose levels or clinical Type-2 diabetes mellitus (T2DM), is associated with symptoms of insomnia, a late chronotype, and shortened sleep duration. These associations have been consistently observed in epidemiological cohort studies and are further supported by genetic analyses^8–11^. Nevertheless, the direct impact of genetic variation on diurnal variation in metabolism remains largely unclear.

Glucose levels fluctuate over the day and are affected by diet, physical activity, stress, sleep, and finally by hormonal factors that maintain physiologically safe levels in circulation^12^. Furthermore, the response to insulin, quantified as insulin sensitivity, is generally higher during the daytime and lower at night. Insulin secretion then typically peaks in the early morning promoting glucose uptake by cells^13^. In contrast, glucagon, which raises blood glucose levels, is more active during periods of fasting, such as overnight. Moreover, during fasting periods the liver releases glucose into the bloodstream and consequently, the liver plays a significant role in maintaining glucose homeostasis^12^.

A noteworthy aspect of T2DM and circulating glucose levels is their unique association with genetic variants from circadian rhythm genes in humans, notably Melatonin Receptor 1B (*MTNR1B*)^14–16^ and Cryptochrome 2 (*CRY2*)^17^. Additionally, both sleep and glucose traits exhibit high heritability and robust genetic associations: Glucose levels are associated with genes expressed in pancreatic tissue, adipose tissue and liver, and the associations have highlighted specific associations for different components of glucose including fasting glucose^14–16^, random glucose^18^, T2DM^19, 20^ or gestational diabetes^19^. Similarly, genome-wide association studies (GWAS) have identified core circadian genes and over 500 other variants associated with chronotype^21^, insomnia^22–24^, and sleep duration^25^. Even larger number of variants have been identified that contribute to biomarkers and metabolites at the normal range^26^. In addition, previous research conducted by our team and others has demonstrated that the disruption of circadian rhythms, such as through night shift work, escalates the risk of cardiovascular diseases and T2DM^27, 28^. Furthermore, insomnia, evening chronotype, and short sleep are established independent risk factors for cardiometabolic diseases. The strength of this connection is further underscored by studies on cardiometabolic traits themselves, revealing that two of the largest effect genetic variants associated with elevated glucose levels and T2DM are located within regulatory regions of *MTNR1B*^14–17^. Previous investigations into *MTNR1B* have established that the risk variant rs10830963 influences insulin secretion, glucose levels. Functional studies on pancreatic islets have revealed that the alternative allele of *MTNR1B* exerts an inhibitory effect on melatonin response, leading to reduced insulin secretion, increased fasting glucose levels, and a heightened risk of T2DM^29^.

These compelling earlier findings prompt the question of whether genetic variants are linked to the diurnal control of glucose levels. In this study, we present evidence using data from 420,000 individuals from the UK Biobank demonstrating that the genetic variants at *MTNR1B* and *CRY2* modulate the diurnal pattern of glucose levels.

## Results

### Glucose levels show time-dependent associations with *CRY2* and *MTNR1B*

While earlier GWAS have revealed a substantial number of variants that contribute to both random and fasting glucose levels^14–16, 19, 20^, it is possible that genetic variants also contribute to timing of these as well. To test a possible diurnal effect from genetic variants to glucose levels, we performed cosinor analysis in glucose measurements and time of sample collection in UK Biobank. Cosinor analysis is a traditional method for examining circadian or diurnal effects that assumes effects are cyclical and time-dependent. We fit sine and cosine models by assuming the time at midnight as time zero. The genome-wide cosine analysis identified two associations, notably *MTNR1B* and *CRY2* (*MTNR1B,* rs10830963, P < 5e-8; *CRY2* rs12419690, P < 5e-8, **Fig. 1A-B**).

**Figure 1.**
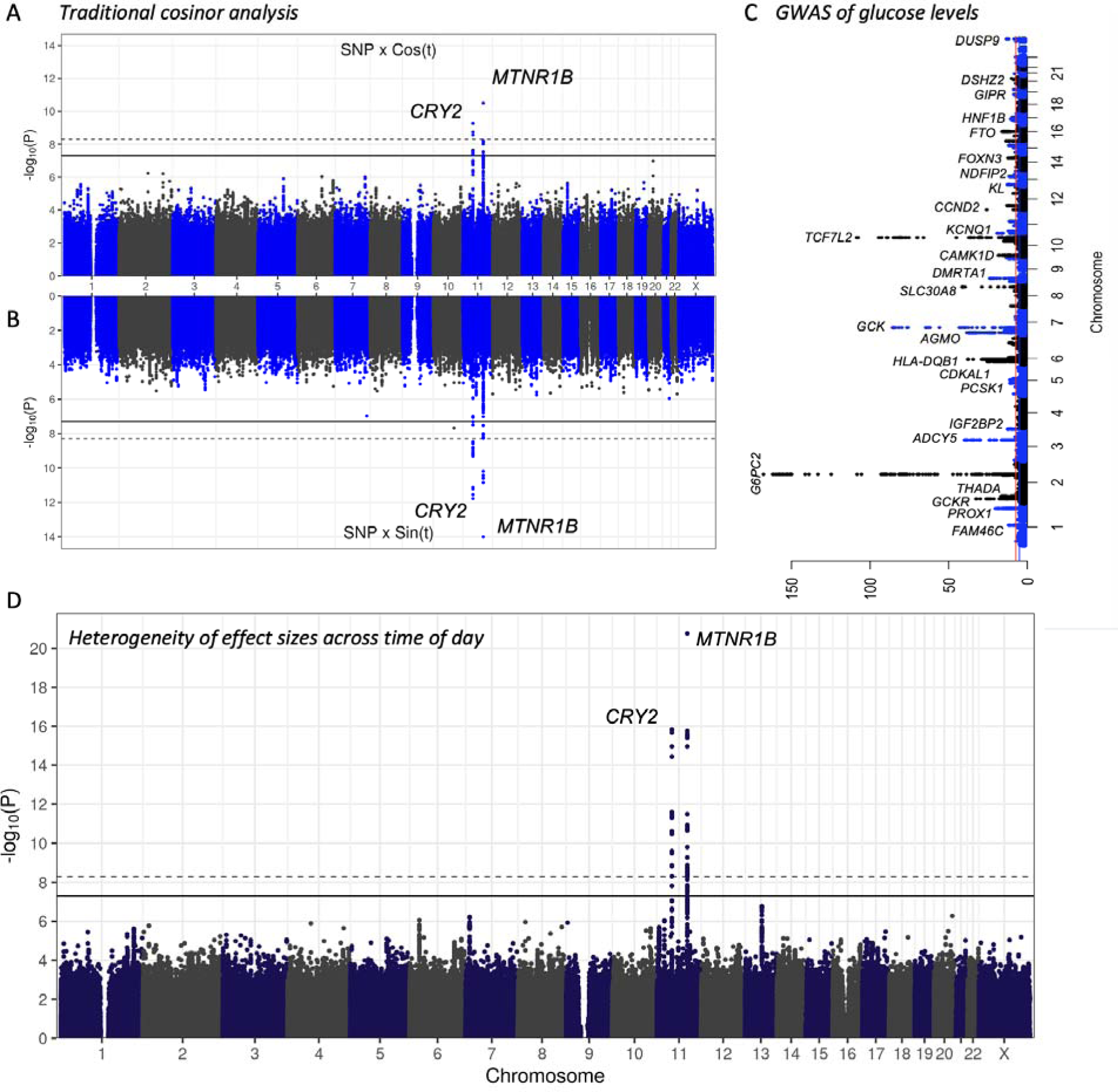
Diurnal association of MTNR1B and CRY2 loci with glucose levels. (A,B) Interaction after accounting for variant main effects shows genome-wide significant interactions over the day at MTNR1B and CRY2 loci in the classical circadian cosinor model. (C) Manhattan plot of GWAS association of glucose levels in UKB shows 66 independent associations. Most significant 25 loci and HLA lead signals are annotated to the closest gene. (D) Estimating heterogeneity of meta-analysis of glucose levels by each hour of the day shows significant heterogeneity at the MTNR1B and CRY2 loci. Solid line represents genome-wide significant threshold P = 5e-8 and dashed line represents P value corrected for multiple testing and number of traits or bins in each analysis.

Curiously, while we discovered 66 genome-wide significant associations with glucose levels (**Fig. 1C, Table S1**), there was no association at either *MTNR1B* or *CRY2* (*MTNR1B* P = 0.27, *CRY2* rs12419690 P = 0.46 **Fig. S1-2**). Thus, we hypothesized that time-dependent effects would also manifest as heterogeneity in association strength throughout the day. This is particularly important as cosinor models rarely fit or perfectly reflect time variation of glucose levels in a population as a result of the daily fluctuation that is modified by meals and hormonal balance. In addition, the vast majority of measurements obtained in most cohorts are collected during daylight hours. Using UK Biobank, we performed GWAS of glucose levels in one hour non-overlapping intervals using measurement times from 9AM to 7PM. Indeed, we detected high heterogeneity at effect sizes only at the *MTNR1B* (rs10830963, P = 1e-22) and *CRY2* loci (rs12419690, P = 1e-16, **Fig. 1D)**.

### Diurnal variants have reversed associations in the morning versus evening

Heterogeneity statistics, while useful for omnibus analysis of variation throughout the day, do not help us understand overall trends between associated time bins. Thus, we visualized the effect sizes used in the heterogeneity analysis to examine overall trends. For *MTNR1B* the effect size for the allele associated with higher glucose levels in the morning showed association with lower glucose levels in the afternoon and evening, supporting a consistent diurnal genetic association with glucose levels (**Fig. 2A, Fig S3-5**). A similar trend was present for *CRY2* (**Fig. S3-5**).

**Figure 2.**
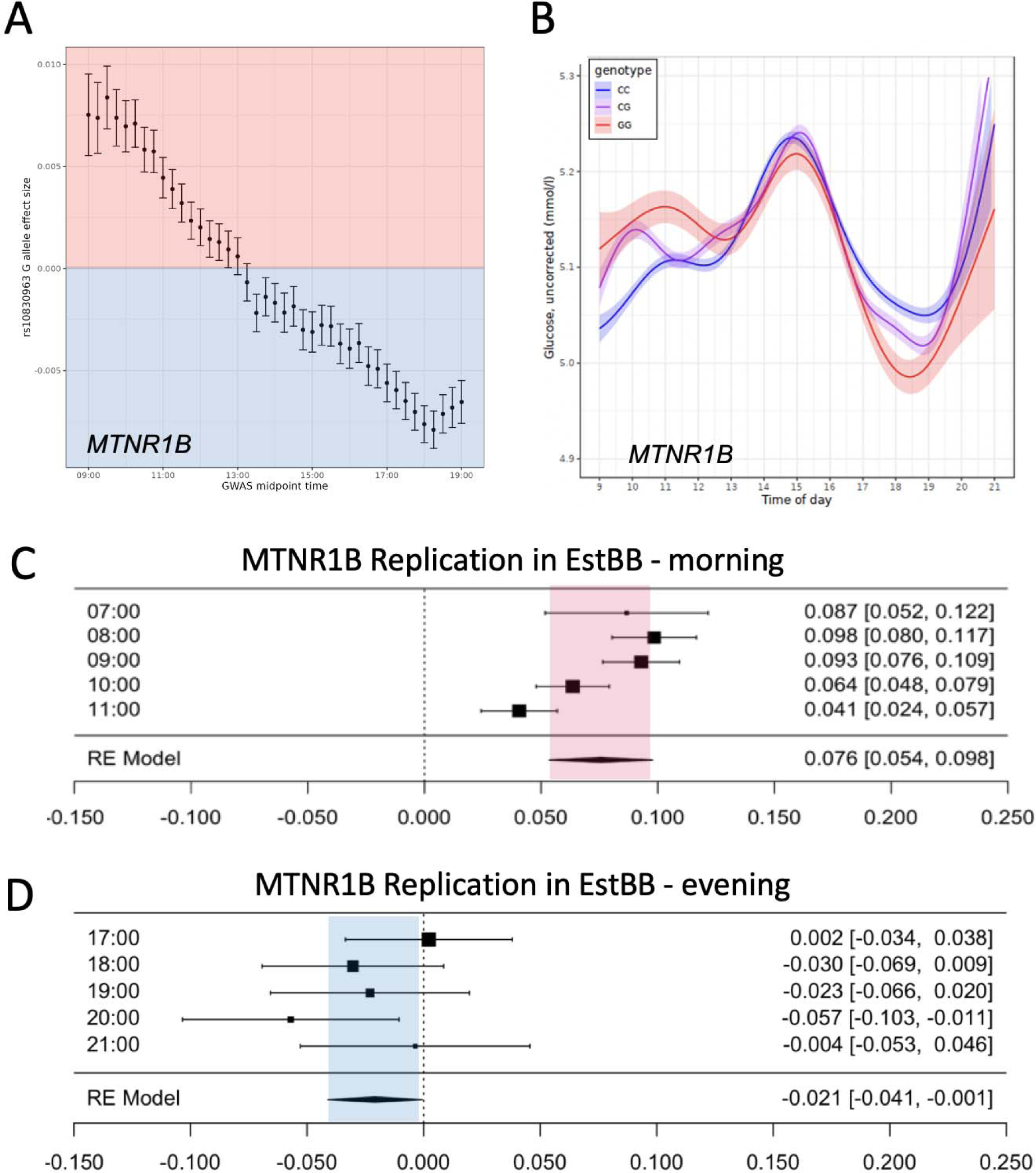
Effect sizes at MTNR1B locus (A) are heterogeneous over the day from 7AM to 7PM showing a decrease in effect that is flipped towards the afternoon and evening. (B) Analysis using generalized linear model (GAM) spline over the day with unadjusted glucose values shows consistent and a higher effect of risk allele rs10830963G in the morning than in the evening. The association of higher effect of risk allele rs10830963G is replicated in Estonian Biobank (C) in the morning and (D) in the evening

Moreover, to formally test the time dependent effect, we fit a spline and tested diurnal effects by genotype using a generalized additive model. We discovered, analogously to the effect size heterogeneity analysis, that risk increasing alleles of *MTNR1B* showed strong intra-day variation in levels, with maximum effect size that was reversed from the morning towards the evening (P < 1e-6, **Fig. 2B**). These findings indicated that genetic variants can show opposite association direction on phenotype depending on time of day when the phenotype was measured. Based on these analyses we estimate that homozygous carriers of the risk alleles have 0.05 mmol/l (*MTNR1B)* and 0.035 mmol/l (*CRY2*) higher glucose levels in the morning.

### Association of *MTNR1B* and *CRY2* with diurnal glucose levels replicates in independent cohorts and with fasting glucose levels

Glucose levels are heavily influenced by environmental factors, most notably meals. We therefore tested the total variance explained by different factors, including time of day, for glucose levels and compared it to other clinical biomarkers measured in the UK Biobank. Ten percent of variation in both fasting and non-fasting levels was explained by time of day, the highest variance explained by time of day across all measured biomarkers^26^.

Given the novel methodology we use in this study, we felt it was important to replicate our findings in an independent second cohort. We therefore tested the associations in the Estonian Biobank (EstBB) in 100,000 individuals. Unlike UK Biobank, EstBB is a clinical collection where laboratory measures are collected as part of hospital admissions and treatment. Both variants replicated, showing similar effects as UKBB, though the heterogeneity of effect sizes across the day at *MTNR1B* (rs10830963 P heterogeneity = 8.79e-28) was substantially more robust than at *CRY2* (rs12419690 P heterogeneity = 0.039, **Fig. S7**). Moreover, the opposite associations from morning to evening were replicated in EstBB for *MTNR1B* rs10830963 (beta morning = 0.076 se = 0.010, P = 1e-13, beta evening = −0.021, se = 0.010 P = 0.03). *CRY2* association was replicated for the morning (beta morning = 0.022, se = 0.004, P = 2e-8) but not in the evening (**Fig. 2C-D**). Overall, the results support a robust association between particularly *MTNR1B* rs10830963 with diurnal glucose levels with opposite effects in the morning versus evening, and across independent cohorts.

As UKB is primarily a non-fasting population, it is possible that the temporal association of *MTNR1B* and *CRY2* with glucose levels would be mediated by eating. To test the possible effect of fasting, we examined the subset of UKB that reported fasting for at least 6, 8 or 12 hours at the time of sample collection (**Fig. S5-6**). Due to substantially smaller sample size of fasting samples (N = 14,000 individuals with 8h or longer fasting), we stratified the analysis by sample collected before 11AM vs. sample collected after 5PM in each fasting category and computed association statistics within the groups. Similar to earlier analysis in the full sample, the effect was in the opposite direction after 5PM (P = 0.001, **Fig. 3A**). Similarly, *CRY2* rs12419690 A allele associated with lower morning glucose levels (P = 0.03) but with higher evening glucose levels (P = 0.001, **Fig. 3B**, and in fasting categories **Fig. S4-5, Table S2**). We observe an association of *MTNR1B* and *CRY2* in all the fasting categories in which we were sufficiently powered, where the *MTNR1B* rs10830963 G allele was associated with higher morning glucose levels (P = 6.9e-5).

**Figure 3.**
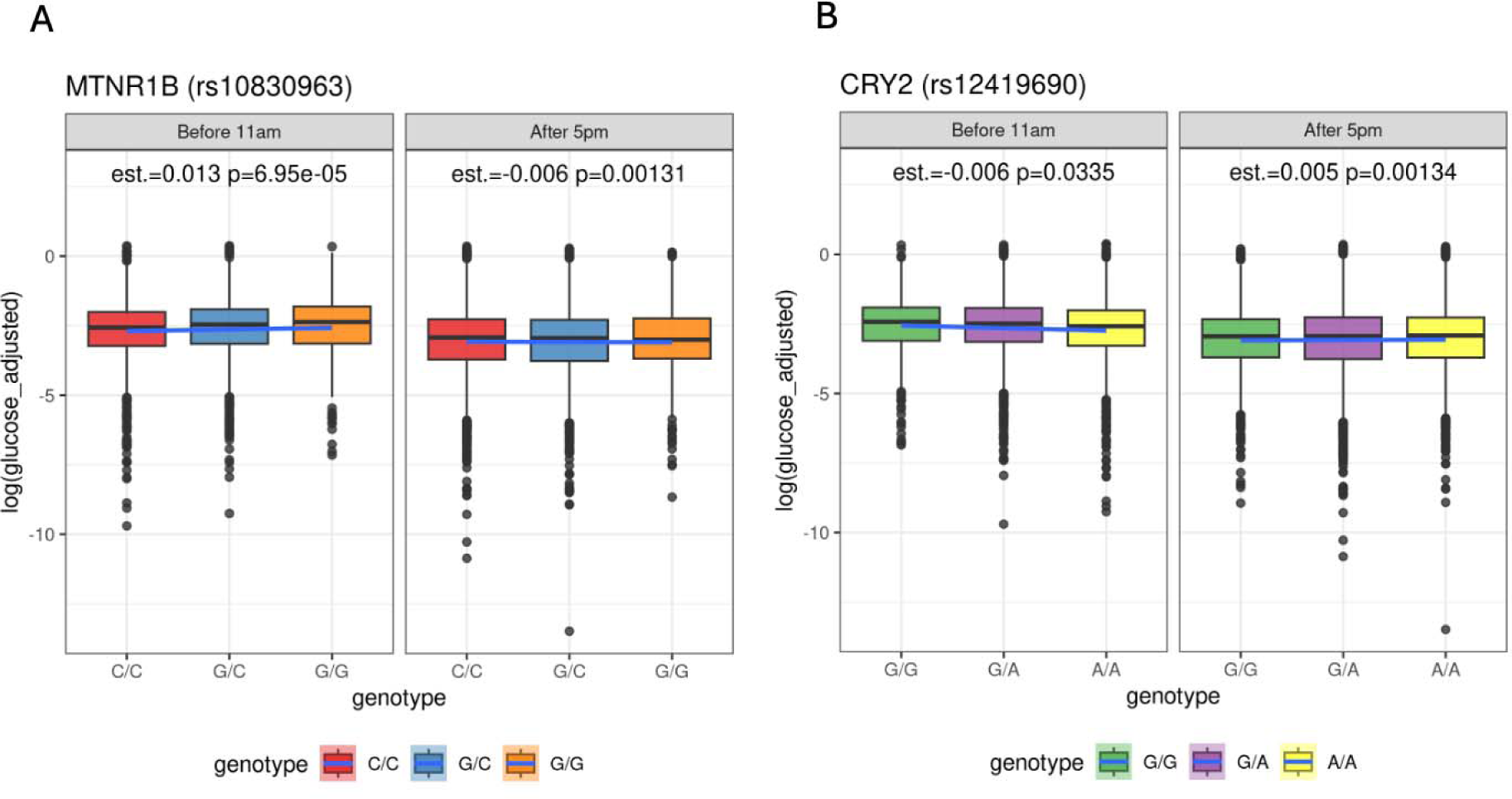
Association of MTNR1B and CRY2 with glucose levels measured in individuals at fasting. Panel A shows MTNR1B rs10830963 and panel B shows data for CRY2 rs12419690. The panels show association with glucose levels in individuals where sample collection individuals in the morning samples (prior to 11AM) or in the evening samples (after 5PM).

### Glucose regulation is connected with sleep and circadian traits

Epidemiological studies and individual genetic associations suggest a tight connection between sleep and glucose traits. Therefore, we tested for a possible shared genetic architecture between glucose and sleep traits using genetic correlation analysis. We saw genetic correlation of fasting glucose levels with napping, insomnia and short sleep, which further connect sleep and circadian traits with glucose homeostasis (**Fig. 4A**). Furthermore, other metabolic traits, such as BMI, associated with insomnia and sleepiness, agreeing with earlier literature^30^.

**Figure 4.**
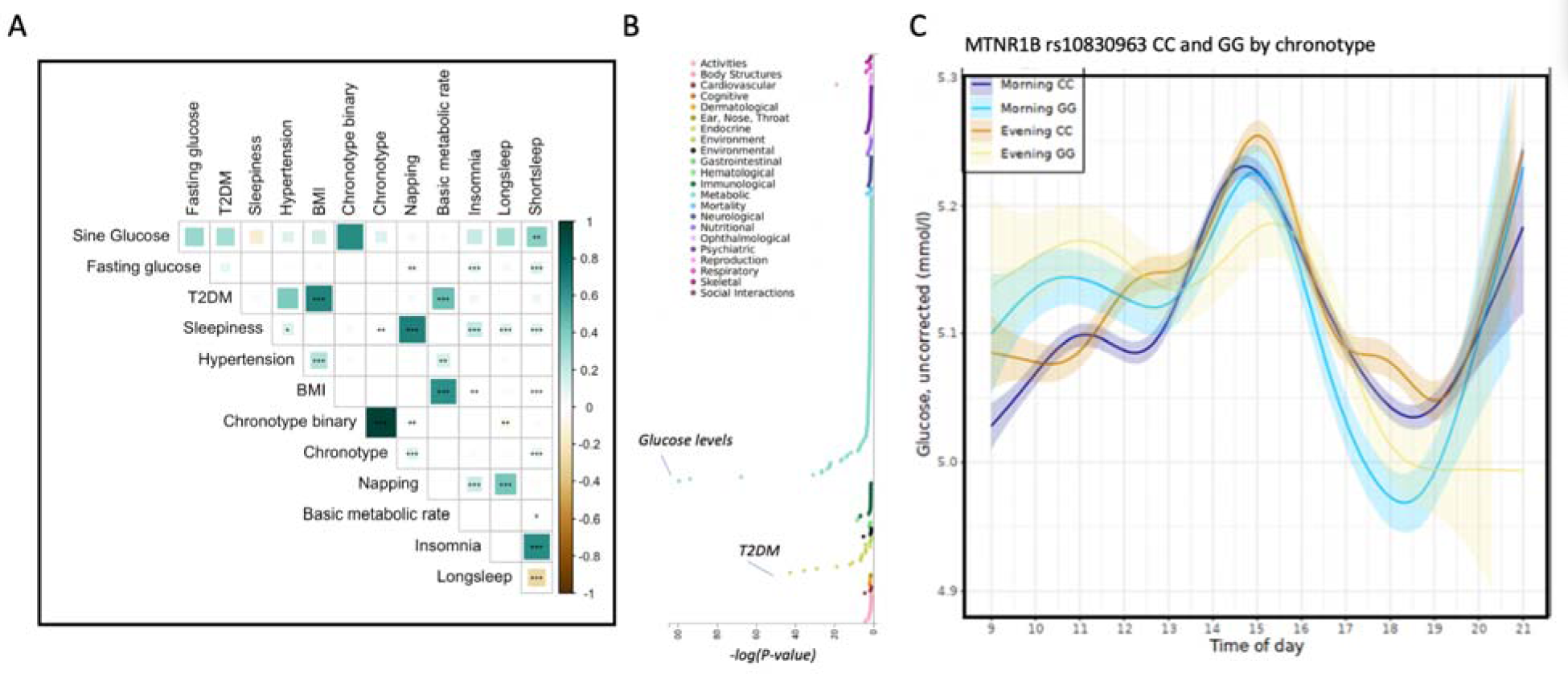
Relationship between sleep and glucose traits. A) Genetic correlation analysis between sleep and circadian traits, and metabolic traits. B) PheWAS of MTNR1B rs10830963 variant shows association with glucose levels and Type-2 diabetes mellitus. C) Glucose levels in individuals with morning versus evening chronotype for MTNR1B rs10830963 variant show consistent effect size independent of chronotype.

We further hypothesized that circadian rhythm coordinates both morningness-eveningness (as measured by chronotype) and diurnal glucose levels. Furthermore, to clarify the phenotypic associations with *MTNR1B* (rs10830963), we performed single-variant-pheWAS, which showed association primarily with metabolic traits and surprisingly not with sleep traits (**Fig. 4B**). We therefore stratified diurnal glucose levels by chronotype, and surprisingly found that MTNR1B risk allele carriers had higher morning glucose levels independently of chronotype (**Fig 4C**). To further understand this finding, we conducted a gene-level-pheWAS of *MTNR1B* and *CRY2*. Our additional analysis revealed independent SNPs at both loci that significantly associate with chronotype but are not in LD (r^2 < 0.2) with the corresponding diurnal glucose associated variants (**Fig. S8**). Therefore, these two loci are known to have multiple regulators, and our analysis suggests that these regulators might independently act in the context of both circadian and diurnal control.

### Daytime sleepiness and napping disrupt the association of *MTNR1B* and *CRY2* with diurnal glucose levels

Existing literature has suggested that sleep disturbance can contribute to higher glucose levels and decreased insulin sensitivity^31^. We therefore tested if chronotype, sleep duration, insomnia, ease of awakening, daytime sleeping or daytime napping modulated the association between diurnal glucose and both the *MTNR1B* and *CRY2* alleles. We observed that *MTNR1B* and *CRY2* variants showed a robust temporal association in individuals who reported insomnia, late chronotype, or short sleep (**Fig. S9-11**). In contrast, individuals who reported daytime napping or sleepiness showed no association between *MTNR1B* or *CRY2* with glucose levels (**Fig. S12**).

In addition, all times of day, we observed a significant epidemiological association between increased glucose levels and insomnia, daytime napping, and daytime sleep (**Fig. S13-15**). We also found a significant association between longer sleep and increased glucose levels (**Fig. S16**) and ease of awakening and decreased glucose levels (**Fig. S17**), but only when glucose was measured in the evening. However, we observed no significant interaction between *MTNR1B* and *CRY2* glucose risk variants and any sleep phenotypes (**Fig. S13-17**), suggesting an independent role of the genetic associations and sleep traits on diurnal glucose variation.

## Discussion

Here, we show that genetic variants at two canonical sleep and circadian genes, *MTNR1B* and *CRY2,* are associated with the timing of glucose levels in humans. In addition, the association we detected was in opposite directions in the morning versus evening, connecting regulation of homeostasis across the day with the regulation of glucose metabolism and with specific genetic loci. The findings indicate that timing of glucose levels is under genetic control. Further, the same genes that contribute to diurnal variation modulate pathology such as T2DM.

One key finding in this paper was the robustness of temporal effects. We observed the association between genetic variants and timing of glucose levels across three methods. A canonical cosinor model tests a sine and cosine of glucose levels by genotype and showed a genome-wide significant association with *MTNR1B* and *CRY2*. We employ multiple non-additive statistical approaches - including classic cosinor analysis from the sleep literature, meta analysis heterogeneity tests, and generalized additive models - to assess the environmental contribution of diurnal variation to genetic effects. This approach robustly identified two known regulators of circadian rhythm and its implications. By integrating alternative modeling strategies into classical GWAS studies, we uncover new biology and can identify additional traits of interest. Applying models which include time-varying components is critical to a comprehensive survey of the genetic architecture of complex traits. Furthermore, it informs how biological time and diurnal rhythms act in context to influence complex traits. Moreover, the benefit of the methods is first the feasibility of applying a cosinor model in GWAS, whereas GAM and heterogeneity analyses can be used in data with even more variable temporal structure as they do not assume a perfect cyclical pattern across the day. These approaches can be used for other time varying data as well, such as circannual variation.

Earlier GWAS on glucose levels have typically discovered *MTNR1B* and *CRY2* associations with fasting glucose levels but not with non-fasting or random glucose levels^15–18^. Our analysis of fasting samples suggest that one of the primary contributing factors for the association most likely is the time component and not the fasting status of the individuals alone. Consequently, including time dependent analysis in GWAS has the potential with larger data sets also to clarify biology bringing additional temporal components into the regulation of homeostasis of metabolites.

It is also intriguing to disentangle the possible mechanisms of how *MTNR1B* and *CRY2* variation modulate glucose levels. Earlier literature on melatonin suggests that even short term melatonin supplementation can induce insulin resistance^29^. We also show in this paper that *MTNR1B* and *CRY2* modulated expression levels to opposite directions in the morning versus in the evening. Melatonin typically indicates biological night, whereas the effects of *CRY2* are potentially related to clock time versus biological time. Imbalance in clock time and biological time is pronounced in shift work or in jet lag - both of which also induce cardiometabolic changes and diseases.

Furthermore, we observe this association independent of insomnia, chronotype or sleep duration. However, sleep problems measured as insomnia or late chronotype had an additional, independent negative effect on glucose levels. Moreover, *CRY2* and *MTNR1B* did not show interaction with sleep traits on glucose levels. First, these findings implicate that sleep has an independent modulatory effect on glucose. Second, genetic variation such as variation at *CRY2* and *MTNR1B* has an independent and additional role of modulating glucose levels in a time dependent fashion.

Overall, the findings indicate that homeostatic glucose levels are modulated by environmental factors such as sleep, as shown here and in previous work. Moreover, inherent genetic factors control the magnitude of overall fasting glucose levels. Finally, genetic factors modulate timing of glucose levels as well. This is likely true in other settings known to regulate these genes, such as the regulation of gestational diabetes in pregnancy by *MTNR1B* and applying time varying models in these unique biological settings will likely reveal additional novel mechanisms of environment dependent regulation. Our findings underscore the diurnal control exerted by *MTNR1B* and *CRY2* on glucose homeostasis, shedding light on the intricate interplay between diurnal rhythms, biological time, metabolism, and fundamental physiological processes.

## Methods

### Cohorts

The UK Biobank is a long-term ongoing study investigating genetic and environmental risk factors that lead to disease. The study contains data on a UK population-based cohort of 503,325 participants aged between 40 and 70. Recruited between 2006 and 2010, participants were invited by means of an invitation letter (UK Biobank Resource 100253) to attend an appointment at one of 22 recruitment centres across the UK. At this baseline visit, participants were asked a series of questions about a number of sociodemographic factors, their lifestyle and their medical history^32^. In addition, physical measurements were taken, cognitive tests performed and biological samples provided (blood, urine and saliva). The 503,325 consenting participants were recruited from ∼9.2 million individuals initially invited: a response rate of 5.47%33. Due to ascertainment bias, UK Biobank participants were more likely to be of higher socioeconomic status (SES) and healthier than the population average in the UK^34^.

The Estonian Biobank is a population-based biobank with 212,955 participants in the current data freeze (2023v1). All biobank participants have signed a broad informed consent form and information on ICD-10 codes is obtained via regular linking with the national Health Insurance Fund and other relevant databases, with the majority of the electronic health records having been collected since 2004^35^.

### Blood and urine sampling

Participants recruited into the UK Biobank study consented to providing a blood sample at their initial (baseline) visit and approximately 480,000 participants gave ∼45ml of blood at the assessment centre. Samples were not asked to fast prior to their visit, but self-report fasting time (data field 74) was recorded. Each participant’s sample tube contained a unique barcode which was scanned on the computerised system, typically a few seconds after sample collection. The timestamp of this barcode scan (data field 3166) was used to determine time of day and date of sample collection, used for deriving the phenotypes and covariates (see below). Part of the blood sample was collected in a serum separator tube, which was then left to clot for 30 minutes at room temperature and then refrigerated before being transported to the central UK Biobank processing facility. Once separated out, the serum was frozen at −80°C for later processing (see UK Biobank Resource 5636; https://biobank.ctsu.ox.ac.uk/crystal/crystal/docs/biomarker_issues.pdf) for 29 serum-based markers.

### Glucose measurements

Blood glucose (mmol/L; data field 30740) was measured (along with other blood biochemistry markers) using the serum samples that were processed at a later date (data field 30741) using the Beckman Coulter AU5800 platform (see resource 1227; https://biobank.ctsu.ox.ac.uk/crystal/crystal/docs/serum_biochemistry.pdf) and extensive quality control and validation was performed to minimise sources of systematic and random variation^36^. We used glucose data for the initial visit only.

### Genotype data

#### UK Biobank

DNA extracted from blood samples provided by UK Biobank participants was genotyped in 106 batches of approximately 4,700 samples per batch. The first 11 batches (N=49,950) were genotyped using a custom Affymetrix genotyping array referred to as the UK BiLEVE^37^. The remaining samples (N=438,427) were genotyped using the very similar (95% overlap) UK Biobank Axiom array, also by Affymetrix (now part of Thermo Fisher Scientific). The arrays contained up to 850,000 probes with ∼650,000 variants used as a dense scaffold for imputation of uncaptured variants, ∼125,000 probes for rare and coding variants, ∼47,000 probes within so-called regions of interest and ∼45,000 markers previously linked to specific phenotypes. Further details of the contents of these arrays can be found at http://tools.thermofisher.com/content/sfs/brochures/uk_axiom_biobank_contentsummary_brochure.pdf. For imputation, markers were included if they had <5% missingness and minor allele frequency (MAF) > 0.0001 and all samples were included except for those with excessive heterozygosity and high missingness. Genotypes were phased with SHAPEIT3 (with 1000 Genomes as a reference panel) and then imputed using both the Haplotype Reference Consortium reference panel^38^ and a combined panel of UK10K and 1000 Genomes phase 3^39^. The two imputed datasets were combined to provide approximately 93 million autosomal variants and nearly 4 million X chromosome variants^40^.

#### Estonian Biobank

All EstBB participants have been genotyped at the Core Genotyping Lab of the Institute of Genomics, University of Tartu, using Illumina Global Screening Array v3.0_EST. Samples were genotyped and PLINK format files were created using Illumina GenomeStudio v2.0.4. Individuals were excluded from the analysis if their call-rate was < 95%, if they were outliers of the absolute value of heterozygosity (> 3SD from the mean) or if sex defined based on heterozygosity of X chromosome did not match sex in phenotype data^41^. Before imputation, variants were filtered by call-rate < 95%, HWE p-value < 1e-4 (autosomal variants only), and minor allele frequency < 1%. Genotyped variant positions were in build 37 and were lifted over to build 38 using Picard. Phasing was performed using the Beagle v5.4 software^42^. Imputation was performed with Beagle v5.4 software (beagle.22Jul22.46e.jar) and default settings. Dataset was split into batches 5,000. A population specific reference panel consisting of 2,695 WGS samples was utilized for imputation and standard Beagle hg38 recombination maps were used. Based on principal component analysis, samples who were not of European ancestry were removed. Duplicate and monozygous twin detection was performed with KING 2.2.7^43^, and one sample was removed out of the pair of duplicates.

### Phenotype derivation

#### UK Biobank

In the UK Biobank cohort, blood glucose levels were taken from data field 30740, which was available in a total of 445,216 participants. To correct the phenotype for sociodemographic and environmental factors, a similar procedure to a previous genetic study of UK Biobank blood traits^26^. To summarise, the raw glucose measure was first log-transformed before the effect of covariates were removed by linear regression. The covariates adjusted for were

- sample dilution factor (factor variable) calculated by quantiling (n=20) data field 30897
- self-report fasting time (factor, data field 74) with those reporting 0 and 1 combined into a single category and those reporting 18 or more combined into another single category
- genotype batch (factor, data field 22000)
- ethnicity (factor, data field 21000) with values 4, 2, 1, 3 and −1 reassigned as 4003, 2004, 1003, 3004 and 6 respectively
- age indicator (factor) set to 50 if age of assessment centre attendance (data field 21003) is 50 or below, set to 78 if age attendance is 78 or above and otherwise set to age at attendance
- age bin (factor, field 21003) set as quintiles of age attended assessment centre (data field 21003)
- BMI indicator (factor) calculated by quantiling (n=50) the result of participant’s weight at attendance in kilograms (data field 21002) divided by the squared height (m) at attendance (data field 50)
- waist-hip ratio corrected for BMI (“WHRadjBMI”, factor) calculated by quantiling (n=50) the residuals from a linear regression of log-scaled hip/waist circumference (i.e. data field 49 divided by data field 48) against BMI (calculated as participant’s weight at attendance in kilograms [data field 21002] divided by squared height (m) at attendance [data field 50])
- centre of attendance (factor, data field 54)
- sex (factor, data field 31)
- month-year indicator (factor) representing a different category for each month and year of attendance (identified from data field 53), with all months of 2006 grouped as a single factor and August to October 2010 as another single factor
- sex-age indicator interaction (using age indicator as defined above)
- sex-fasting time interaction (using fasting time as defined above)
- sex-ethnicity interaction (using ethnicity as defined above)
- sex-BMI indicator interaction (using BMI indicator as defined above)
- sex-WHRadjBMI interaction (using WHRadjBMI as defined above)
- age bin-fasting time interaction (using fasting time as defined above)
- glucose assay date (factor, data field 30741)
- glucose aliquot (factor, data field 30742)

The residuals from this large regression were used as the adjusted (log-scale) glucose measurement. The inverse-normalised phenotype was generated by rank-normalising these residuals in European-ancestry participants only.

#### Estonian Biobank

In the Estonian Biobank cohort, the glucose measurement values and timepoints were obtained from electronic health records (LOINC laboratory test code 14749-6 “Glucose in Serum or Plasma”). For each participant, we opted to use the earliest possible measurement. Participants with values below <1 and above >15 were excluded from the study.

### Genome-wide Association Analyses

To assess whether genetic variants have a time-dependent effect on glucose levels, we performed two types of genome-wide association analyses in the UK Biobank, which utilised the blood sample collection time.

### Sinusoidal analysis

For primary analyses, we calculated genetic associations by analysing the residualised glucose phenotype using REGENIE v3.1.1^44^ with a sinusoidal GxE interaction term (REGENIE step 2 “--interaction” flag), where the interaction term was defined as either *sin* (2*πt*) or *cos* (2*πt*) where *t* is the proportion of the day passed (since midnight) at blood draw time, calculated as

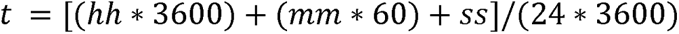

and with hh:mm:ss being the timestamp for blood sample collection, extracted from data field 3166. Ideally, both sine and cosine interaction terms would be included in the same REGENIE association test model, as they capture effects that are out of phase with one another and so together can capture time-of-day-interaction effects across the entire day. REGENIE only allows a single interaction effect, so the sine and cosine interaction terms were each assessed in separate REGENIE runs. For these analyses, we report the “ADD-INT_SNP” test results output by REGENIE, which represent the additive effect of the variant’s on the phenotype as a function of the interaction term, corrected for the variant’s additive effect independent of the interaction term.

In both the sine and cosine interaction analyses, we adjusted for the categorical covariates sex (data field 31), genotyping array (3 categories derived from data field 22000; UKBiLEVE [batches −11 to −1], UKB Axiom initial release [batches 1 to 22] and UKB Axiom full release [batches 23 to 95]), centre of attendance (data field 54), self-report fasting time (derived from data field 74 as described above) and continuous covariates age at assessment (data field 21003) and genetic principal components 1 to 10 (data field 22009).

These analyses primarily identify time-dependent additive associations, which best capture the effect of variants whose effect allele increases the phenotype when it is higher than the mean level, reduces the phenotype when it is lower than the mean level and has little-to-no effect when the phenotype is near average levels. Figure Y shows a schematic representation of an imaginary variant that has no purely additive effect on the phenotype, but has a time-dependent additive effect. Note that in this idealised scenario a standard GWA analysis that does not correct for time of day would show no significant association for this variant.

### Windowed analysis

As a sensitivity analysis, we used an independent method to validate findings from the sinusoidal interaction genetic analyses. To do this, we split individuals into 6 groups based on their time of day of blood draw: [08:00,10:00), [10:00,12:00), [12:00, 14:00), [14:00, 16:00), [16:00, 18:00) and [18:00, 20:00). We then performed a genetic association analysis of the residualised glucose levels for each group independently using REGENIE v3.1.1 and correcting for the same covariates as in the sine and cosine analyses. We then meta-analysed these results.

### Estonian Biobank

For the replication analyses, we divided the Estonian Biobank measurements into 24 groups according to the measurement time point hours. Association analyses in Estonian Biobank were carried out for all variants with an INFO score > 0.4 using the additive model as implemented in REGENIE v3.2 with standard quantitative trait settings^44^. Linear regression was carried out with adjustment for current age, age², sex and 10 PCs as covariates, analyzing only variants with a minimum minor allele count of 2.

### Analysis of *MTNR1B* and *CRY2* risk variants in fasting individuals

To investigate the role of food consumption mediating the association between *MTNR1B* and *CRY2* risk variants and glucose levels, we subset the individuals in the UKB with glucose measurements to those with reported fasting times of 6, 8, or 12 hours at the time of sample collection. Due to substantially smaller sample size of fasting samples, we grouped individuals whose samples were collected before 11am and those with sample collection after 5pm. For each time point (before 11am, after 5pm) and risk variant (*MTNR1B, CRY2*), we fit a linear regression for adjusted glucose (derived as described in methods, phenotype derivation) with risk genotype and sex, age at baseline, array, center, and first ten principal components of genetic variance as covariates.

### Analysis of sleep and circadian traits and *MTNR1B*, *CRY2* risk variants

To assess the role of sleep phenotypes in interaction with the risk variants at *MTNR1B* and *CRY2* loci, we assessed questionnaire data from the UK biobank for sleep duration (Field ID 1160), insomnia (Field ID 1200), chronotype (Field ID 1180), napping during the day (Field ID 1190), daytime dozing or sleeping (Field ID 1220), and ease of awakening in the morning (Field ID 1170). For each time point (before 11am, after 5pm) and risk variant (*MTNR1B, CRY2*), we fit a linear regression for adjusted glucose (previously described) with risk variant, sleep phenotype, interaction between sleep phenotype and risk variant, and sex, age at baseline, array, center, and first ten principal components of genetic variance as covariates.

## Supporting information

Table S1 (glucose association statistics)

## Supplementary Figures and Tables

**Supplementary Figure 1.**
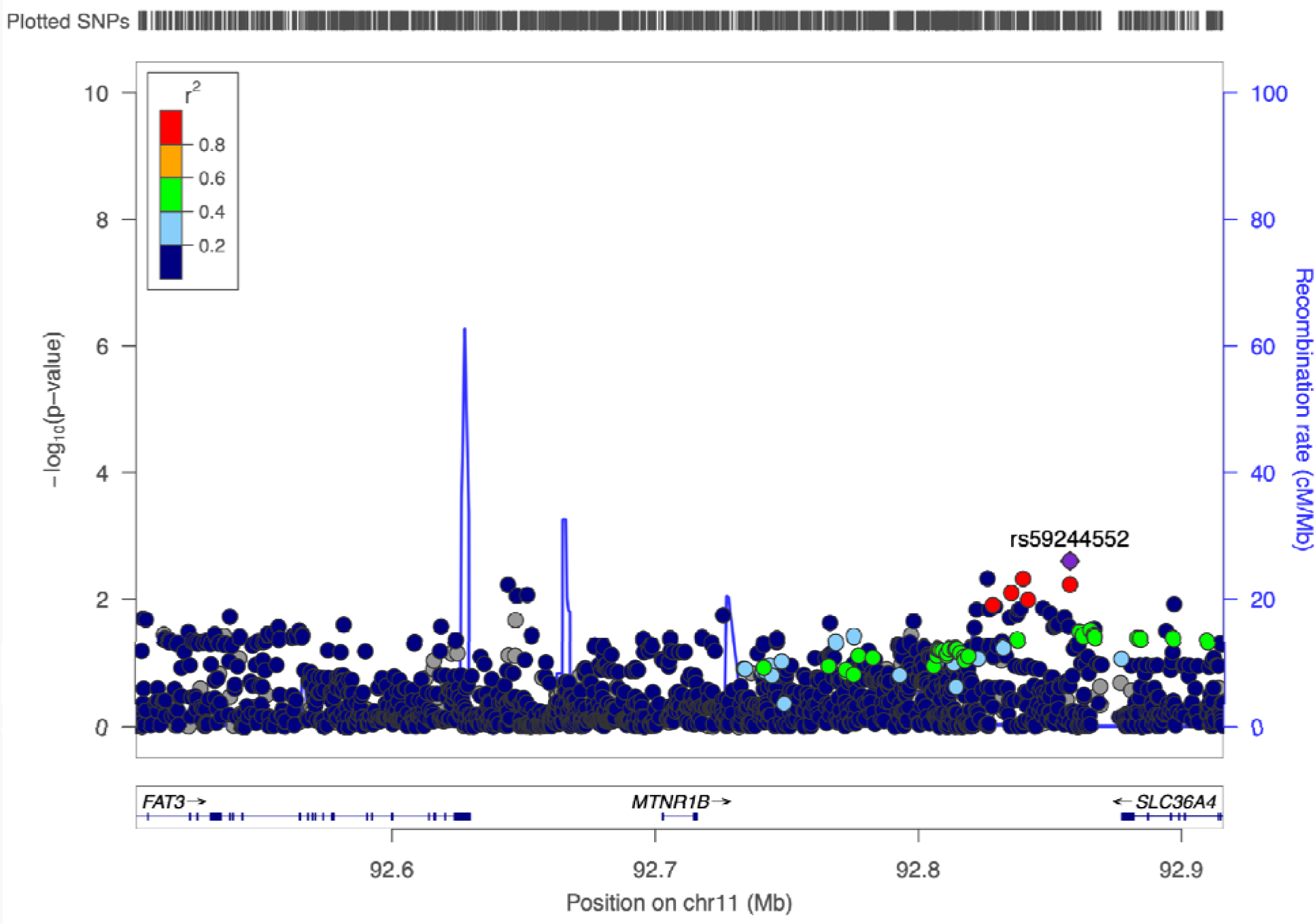
Association of variants at the *MTNR1B* locus with glucose levels. We computed genome-wide association statistics for glucose levels (field id = 30740) in the UKBB. The regional analysis did not show significant association MTNR1B rs10830963 MTNR1B (rs10830963 P = 0.27) or other variants at the locus.

**Supplementary Figure 2.**
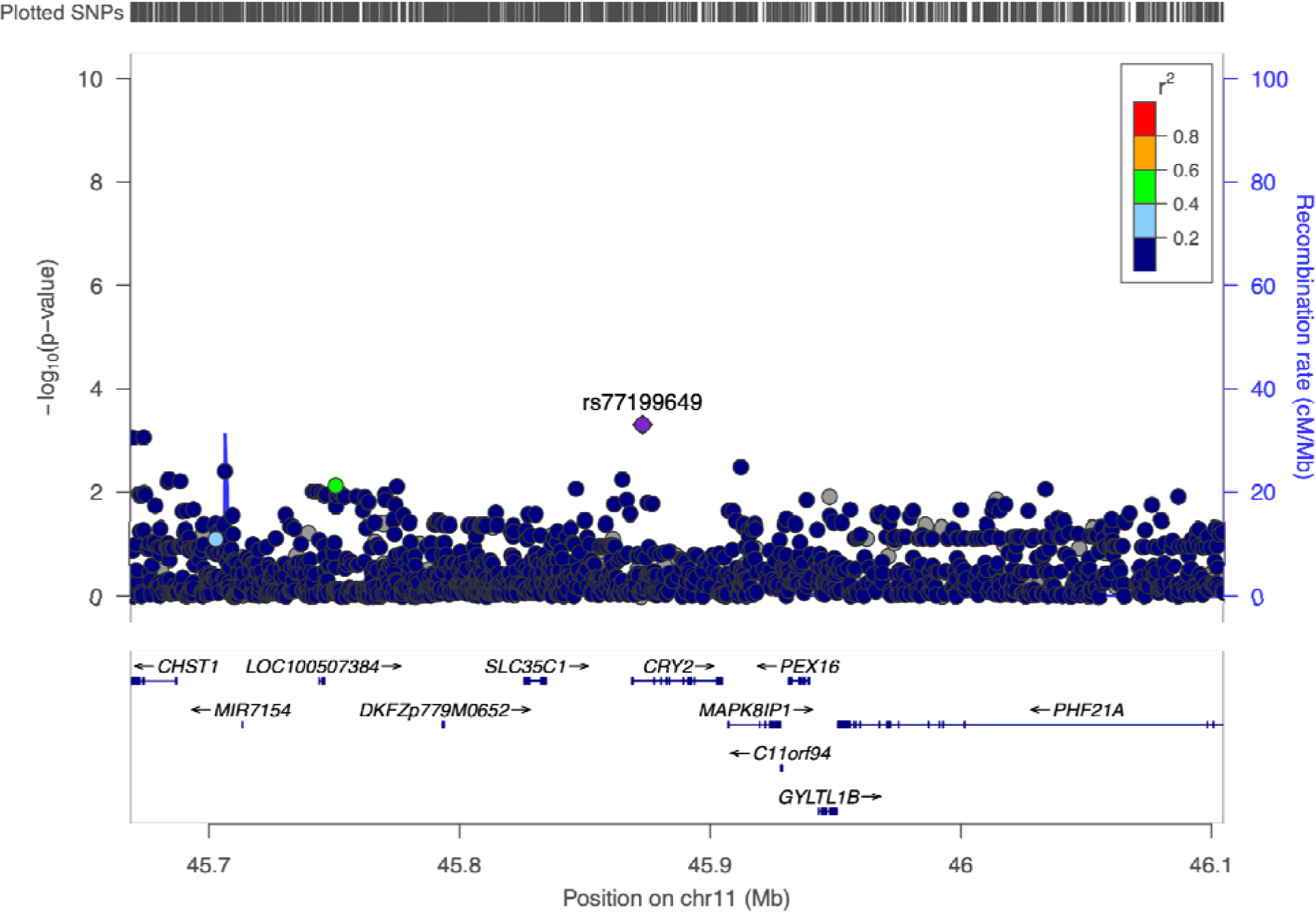
Association of variants at the *CRY2* locus with glucose levels. We computed genome-wide association statistics for glucose levels (field id = 30740) in the UKBB. The regional analysis did not show significant association with CRY2 variant (rs12419690 P = 0.46) or other variants at the locus.

**Supplementary Figure 3.**
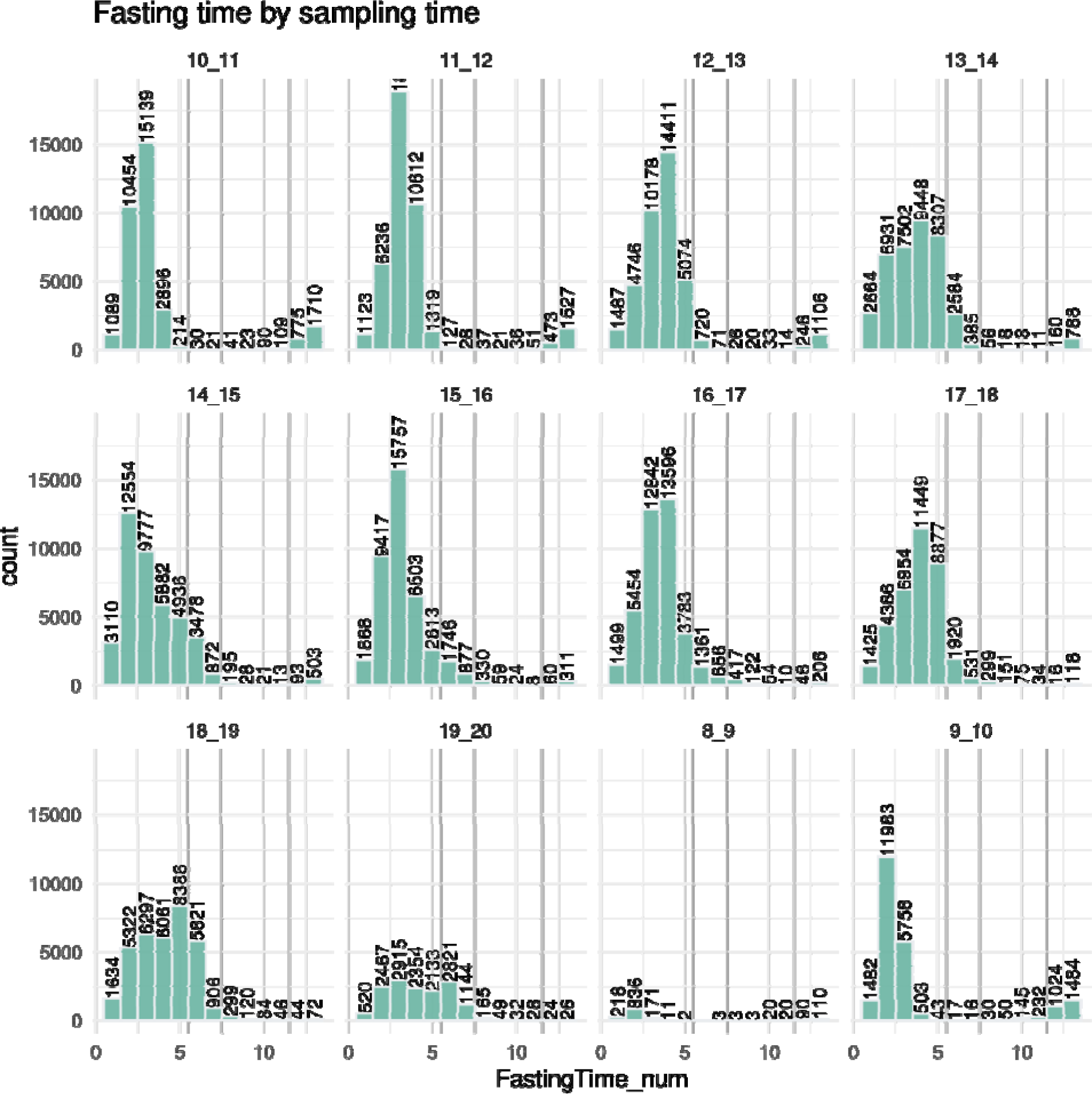
Fasting time stratified by time of sample collection. We binned data by time of measurement from 8AM to 8PM, and by number of hours fasting. We then computed the number of individuals in each fasting bin by hours of fasting.

**Supplementary Figure 4.**
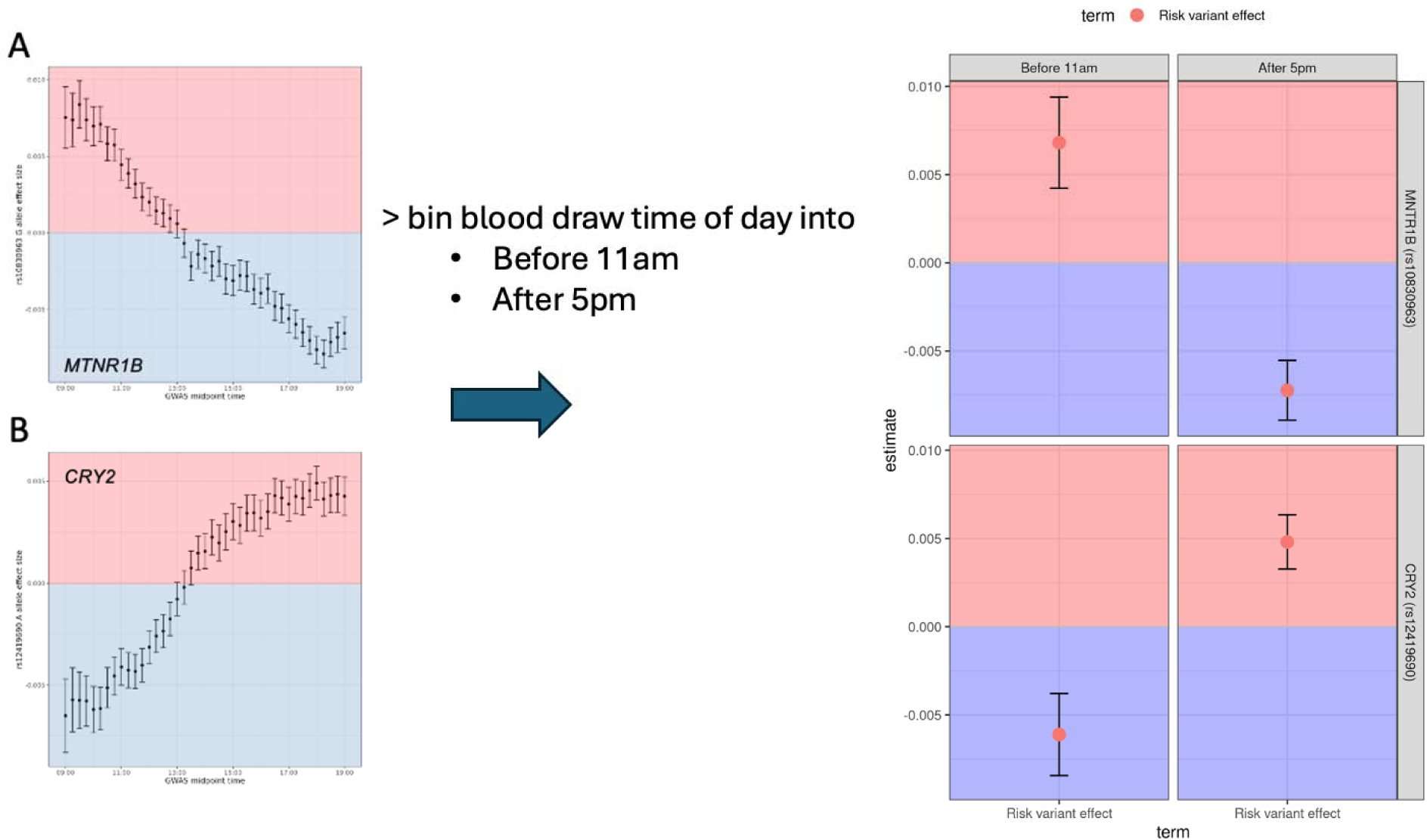
Binning process for morning and evening stratification. Due to the substantially smaller sample size of individuals fasting in UKBB, we stratified the analysis by sample collected before 11am vs sample collected after 5pm.

**Supplementary Table 2.** Number of fasting samples at each blood collection time and fasting time.

| Fasting time | Blood collection time | n samples |
| --- | --- | --- |
| 6 hours or more | Before 11am | 12,309 |
| 6 hours or more | After 5pm | 31,441 |
| 8 hours or more | Before 11am | 12,133 |
| 8 hours or more | After 5pm | 3,539 |
| 12 hours or more | Before 11am | 10,573 |
| 12 hours or more | After 5pm | 607 |

**Supplementary Figure 5.**
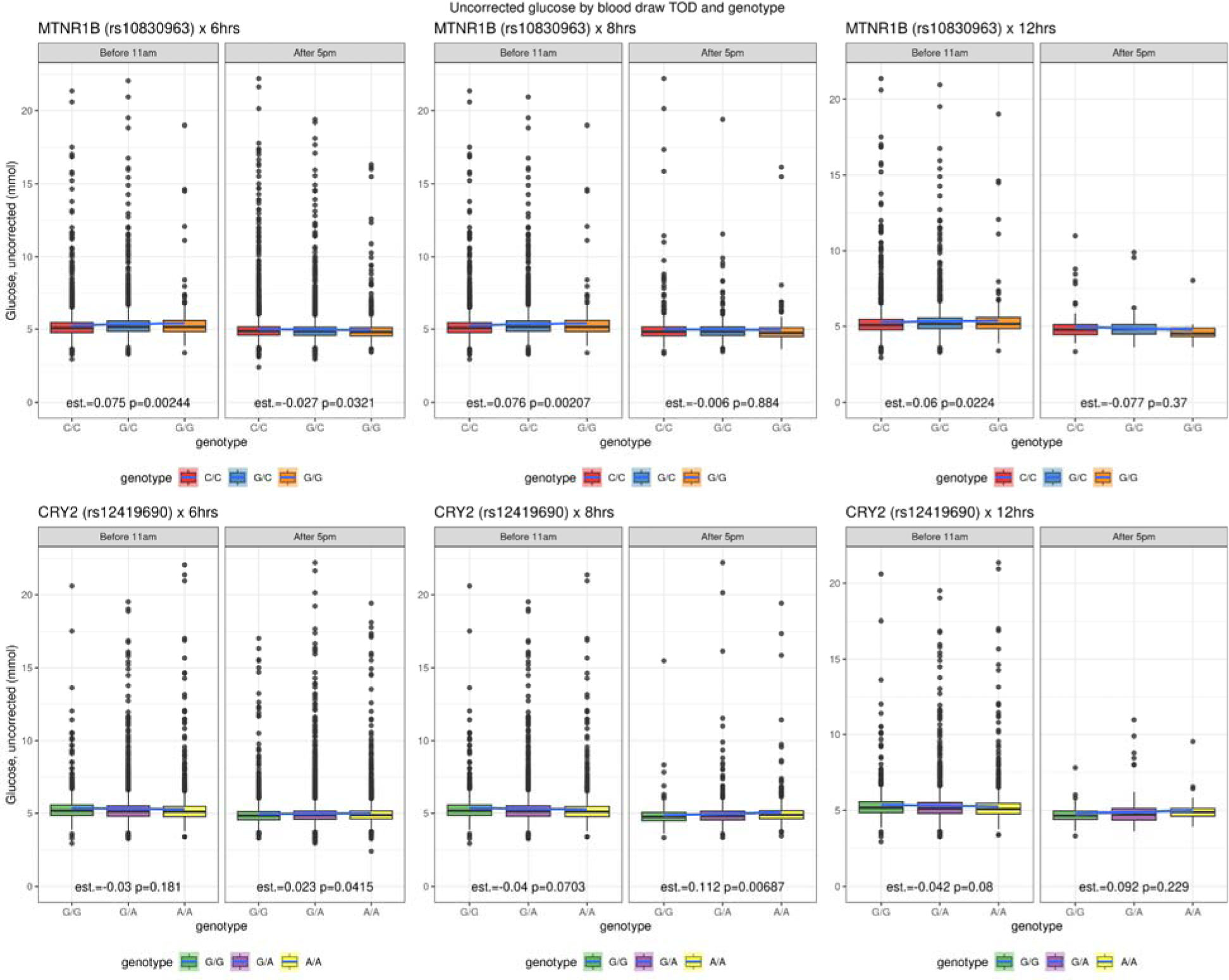
Comparison distribution of association in the morning and evening. Pairwise comparison of reference and alternative risk alleles in fasting individuals with sample collection time before 11AM or after 5PM with unadjusted glucose levels.

**Supplementary Figure 6.**
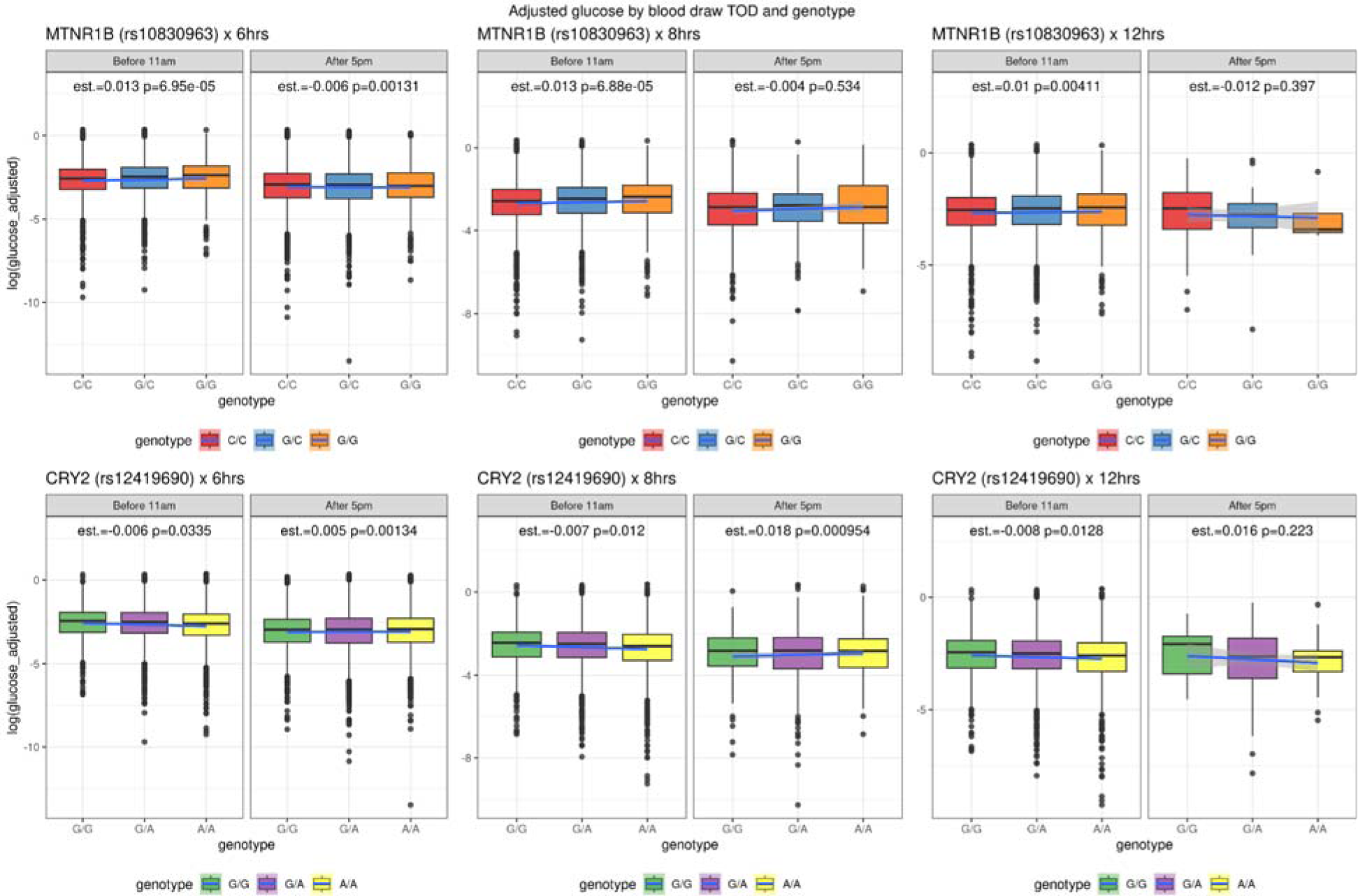
Comparison distribution of association of the residual glucose levels in the morning and evening. Comparison of reference and alternative risk alleles in fasting individuals with sample collection time before 11AM or after 5PM with the residual glucose levels (see methods).

**Supplementary Figure 7.**
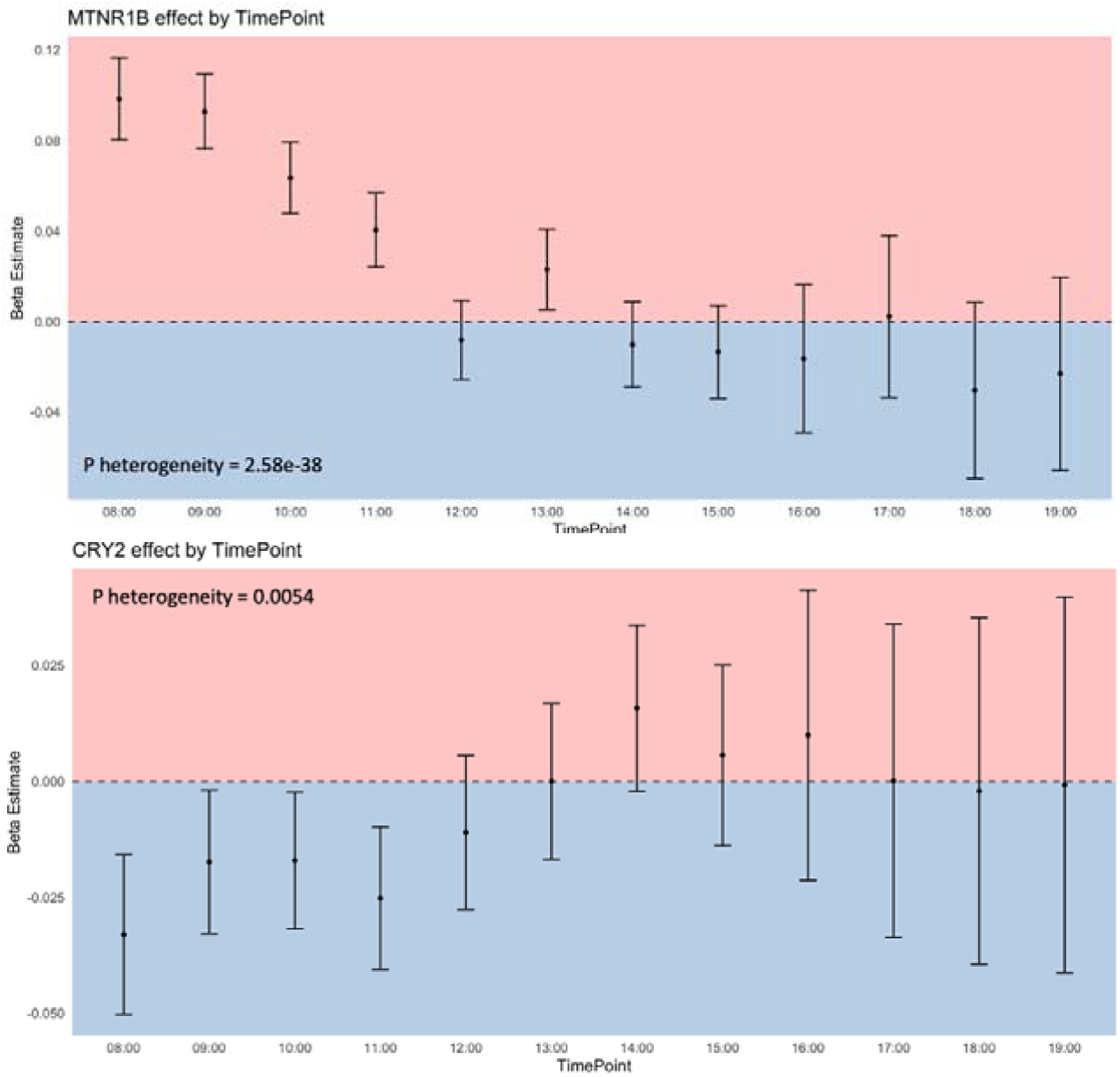
Effect size distribution of *MTNR1B* and *CRY* in Estonian Biobank. We computed association statistics for each hour of the day in Estonian Biobank and visualize the effect estimate for effect allele stratified by measurement time.

**Supplementary Figure 8.**
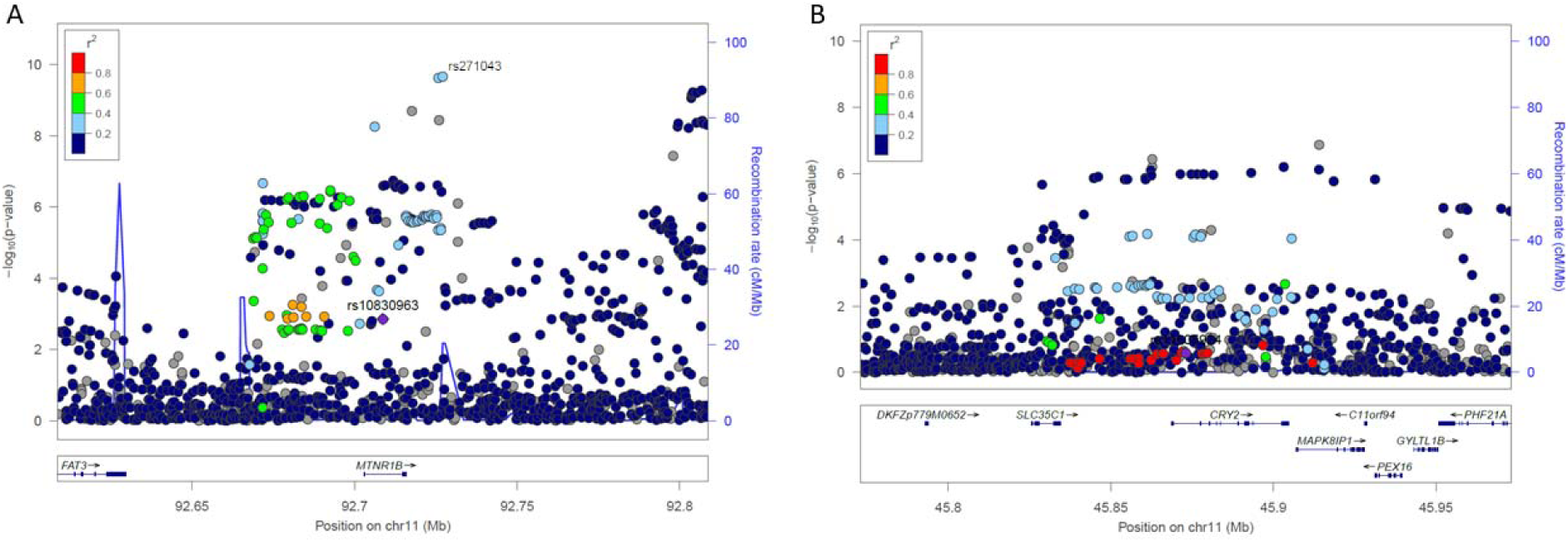
Association of *MTNR1B* and *CRY2* loci with chronotype. Regional association plot for chronotype at the A) *MTNR1B* and B) *CRY2*. We observed association both at *MTNR1B* and at *CRY2* locus with chronotype. However, the variants that associate with chronotype are not in strong LD (color scale) with the variants that associate with glucose levels.

**Supplementary Figure 9.**
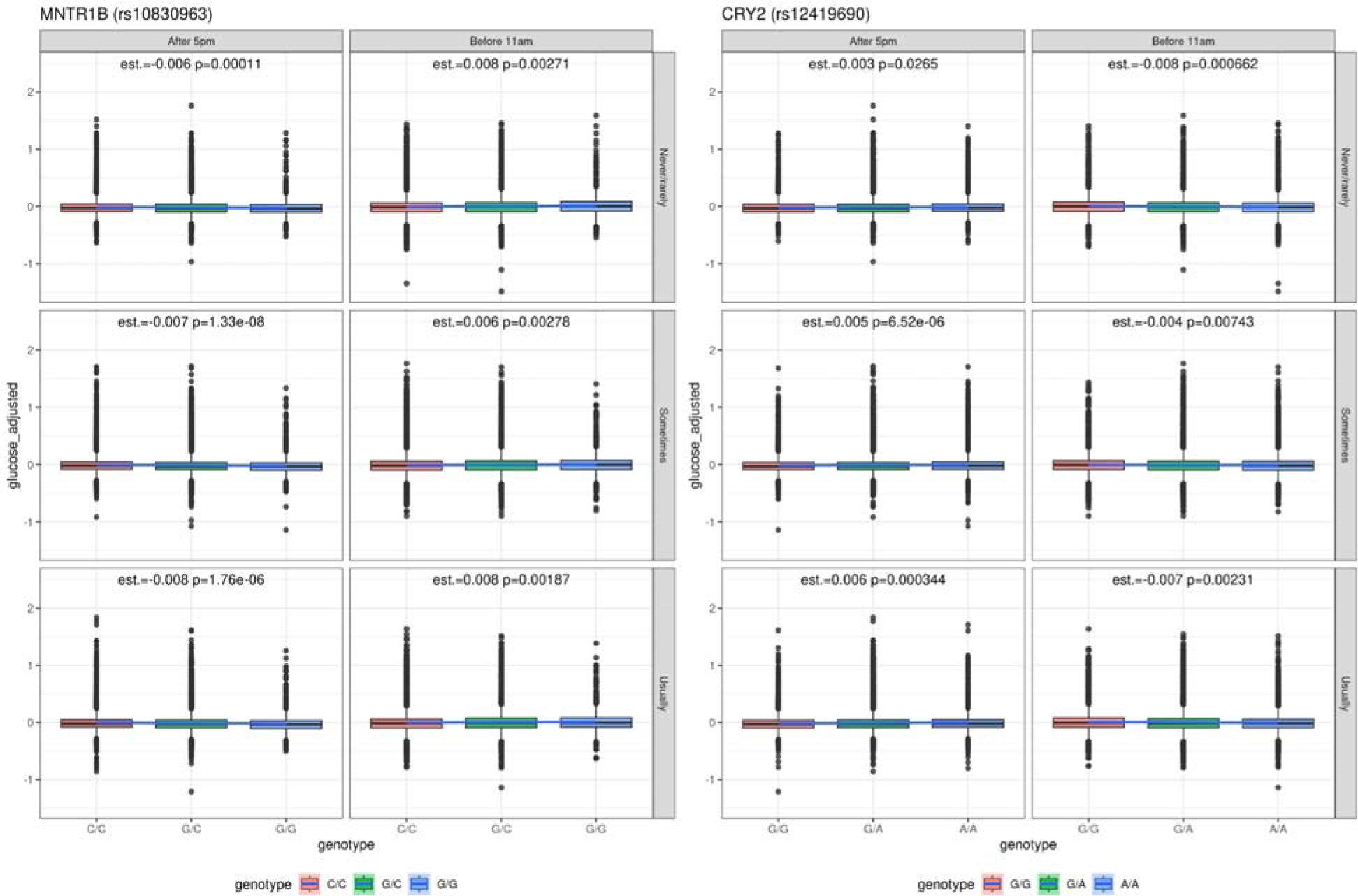
Comparison distribution of association of the residual glucose levels in the morning and evening stratified by insomnia. We computed the effect size for *CRY2* and *MTNR1B* effect alleles, stratified by time of day of sampling and insomnia. The estimate reflects additive genotype effect and p-value.

**Supplementary Figure 10.**
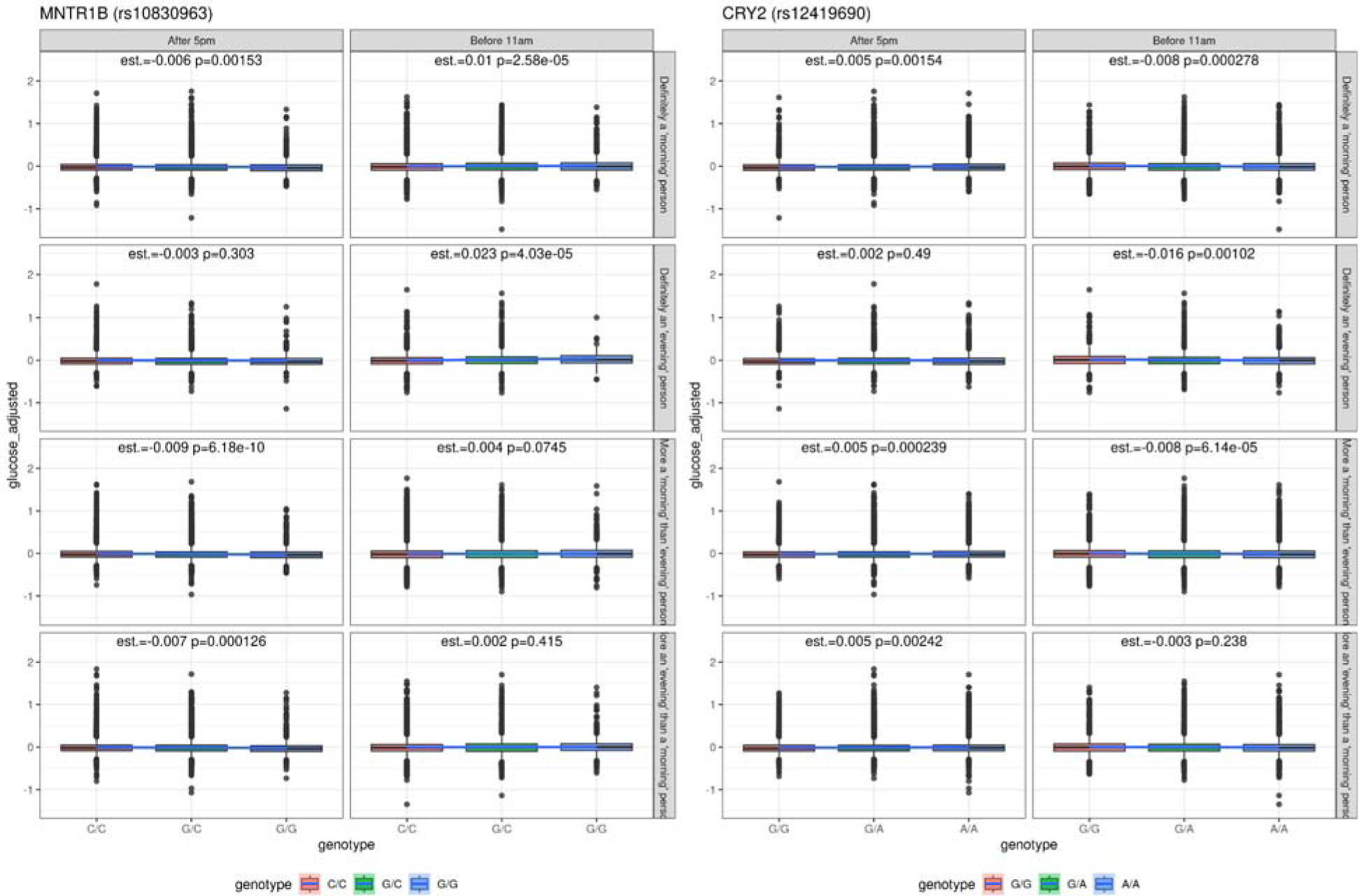
Comparison distribution of association of the residual glucose levels in the morning and evening stratified by chronotype. Effect of *CRY2* and *MTNR1B* risk variants, stratified by time of day of sampling and chronotype. The estimate reflects additive genotype effect and p-value.

**Supplementary Figure 11.**
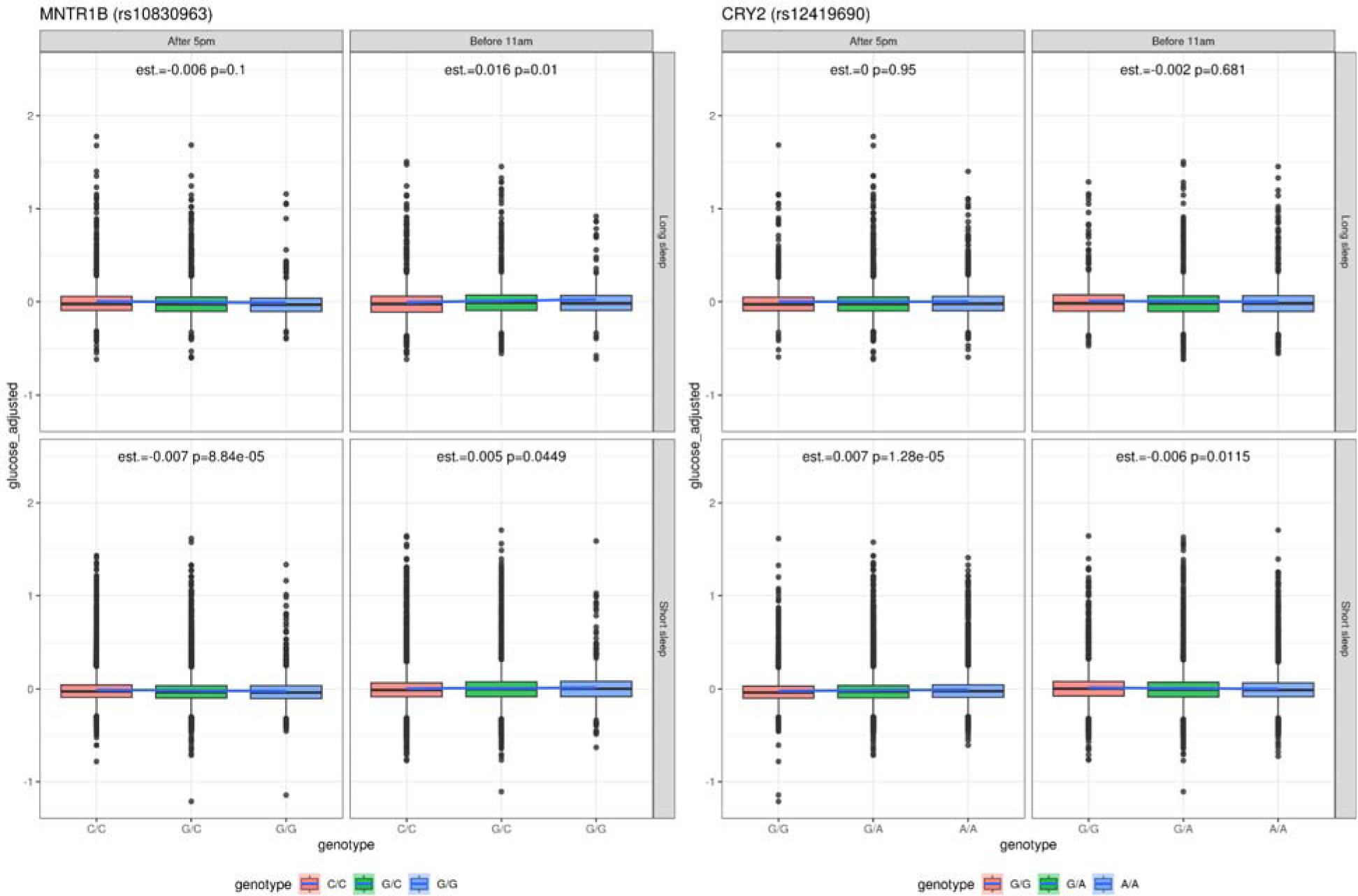
Comparison distribution of association of the residual glucose levels in the morning and evening stratified by sleep duration. Effect of *CRY2* and *MTNR1B* risk variants, stratified by time of day of sampling and sleep duration. Short sleep was defined as 6 hours or less, and long sleep 9 hours or more. The estimate reflects additive genotype effect and p-value.

**Supplementary Figure 12.**
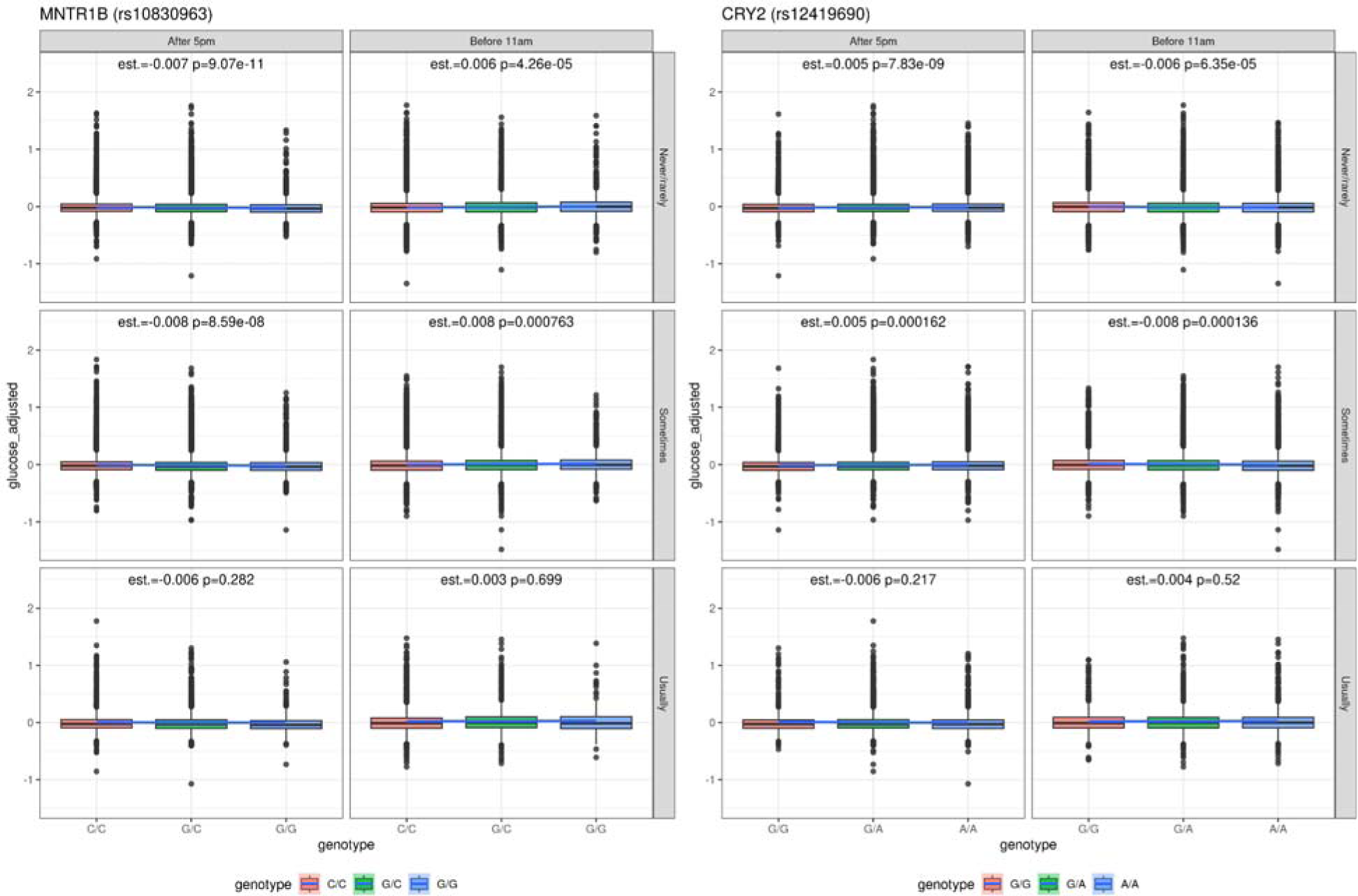
Comparison distribution of association of the residual glucose levels in the morning and evening stratified by daytime napping/sleeping. Effect of *CRY2* and *MTNR1B* risk variants, stratified by time of day of sampling and daytime napping/sleeping. The estimate reflects additive genotype effect and p-value.

**Supplementary Figure 13.**
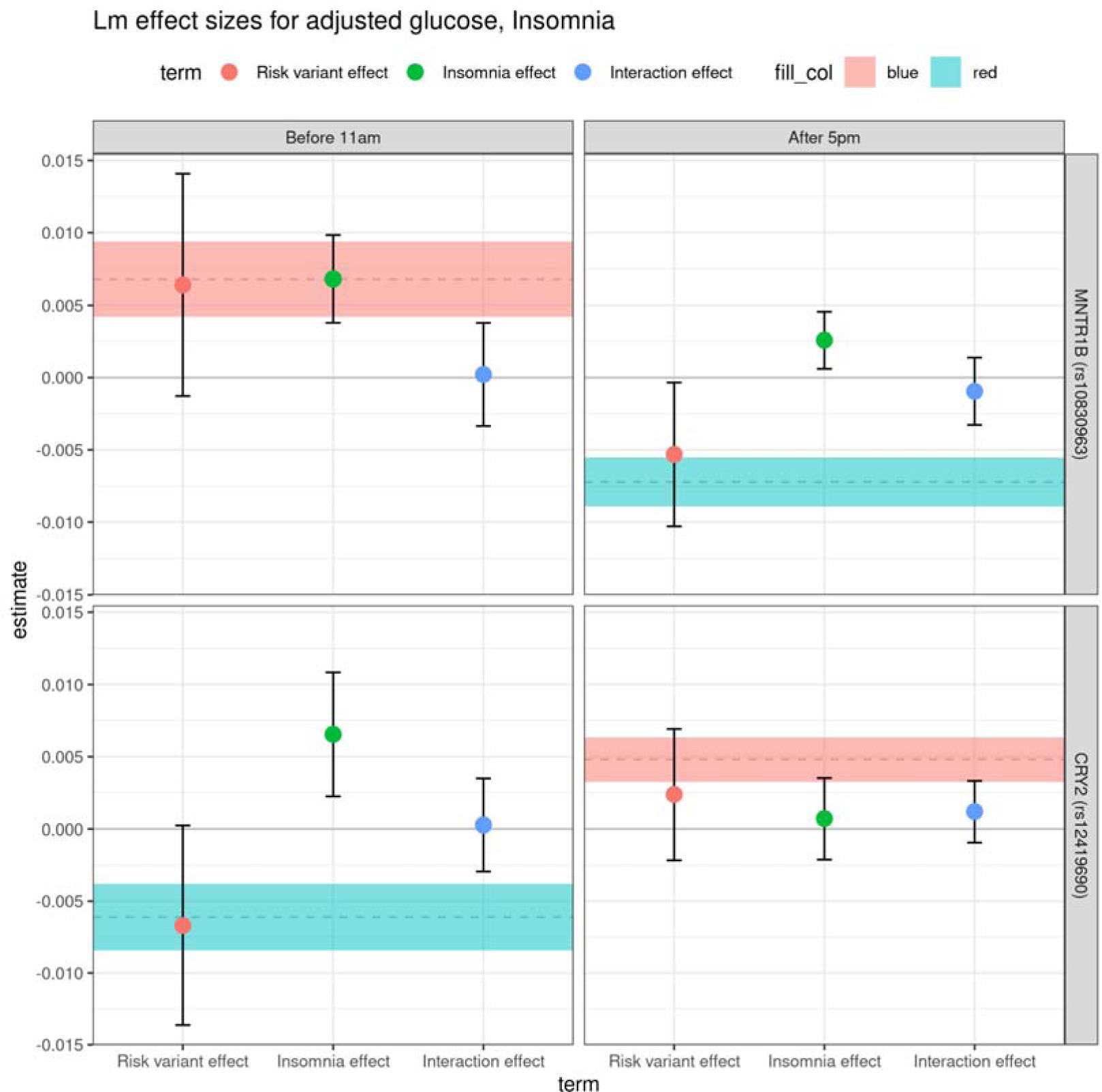
Effect of insomnia, risk genotype and interaction between insomnia and genotype on glucose levels. We computed the association of insomnia (green), risk variant (red) and interaction effect (blue) on glucose levels. This shows an effect of insomnia on glucose both in the morning and evening but no interaction between the genotype and insomnia effect. Dashed lines and shaded area in background represent risk variant effect size and 95% confidence intervals over all samples.

**Supplementary Figure 14.**
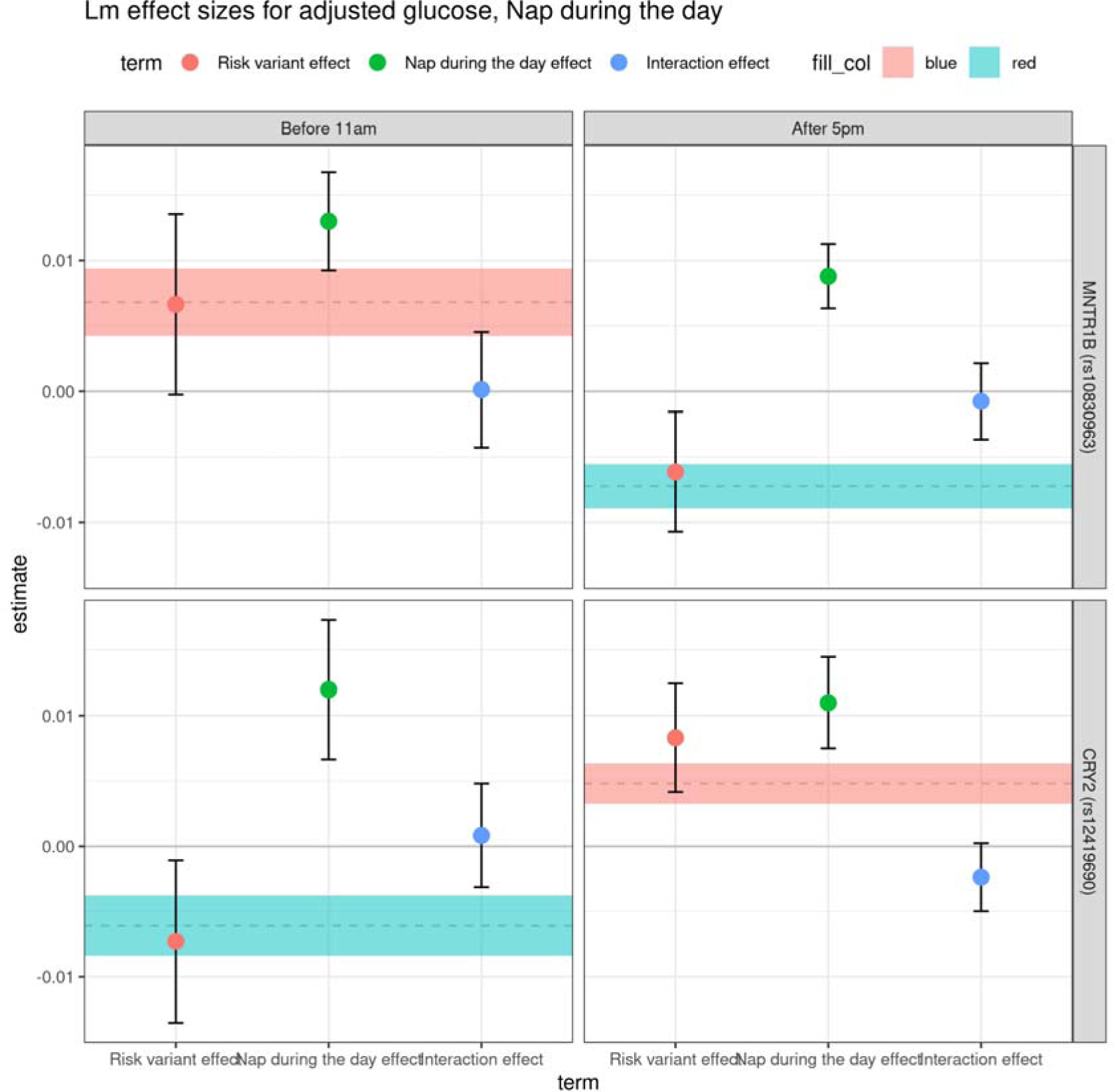
Effect of napping, risk genotype and interaction between insomnia and genotype on glucose levels. We computed the association of nap during the day (green), risk variant (red) and interaction effect (blue) on glucose levels. This shows an effect of napping on glucose both in the morning and evening but no interaction between the genotype and napping effect. Dashed lines and shaded area in background represent risk variant effect size and 95% confidence intervals over all samples.

**Supplementary Figure 15.**
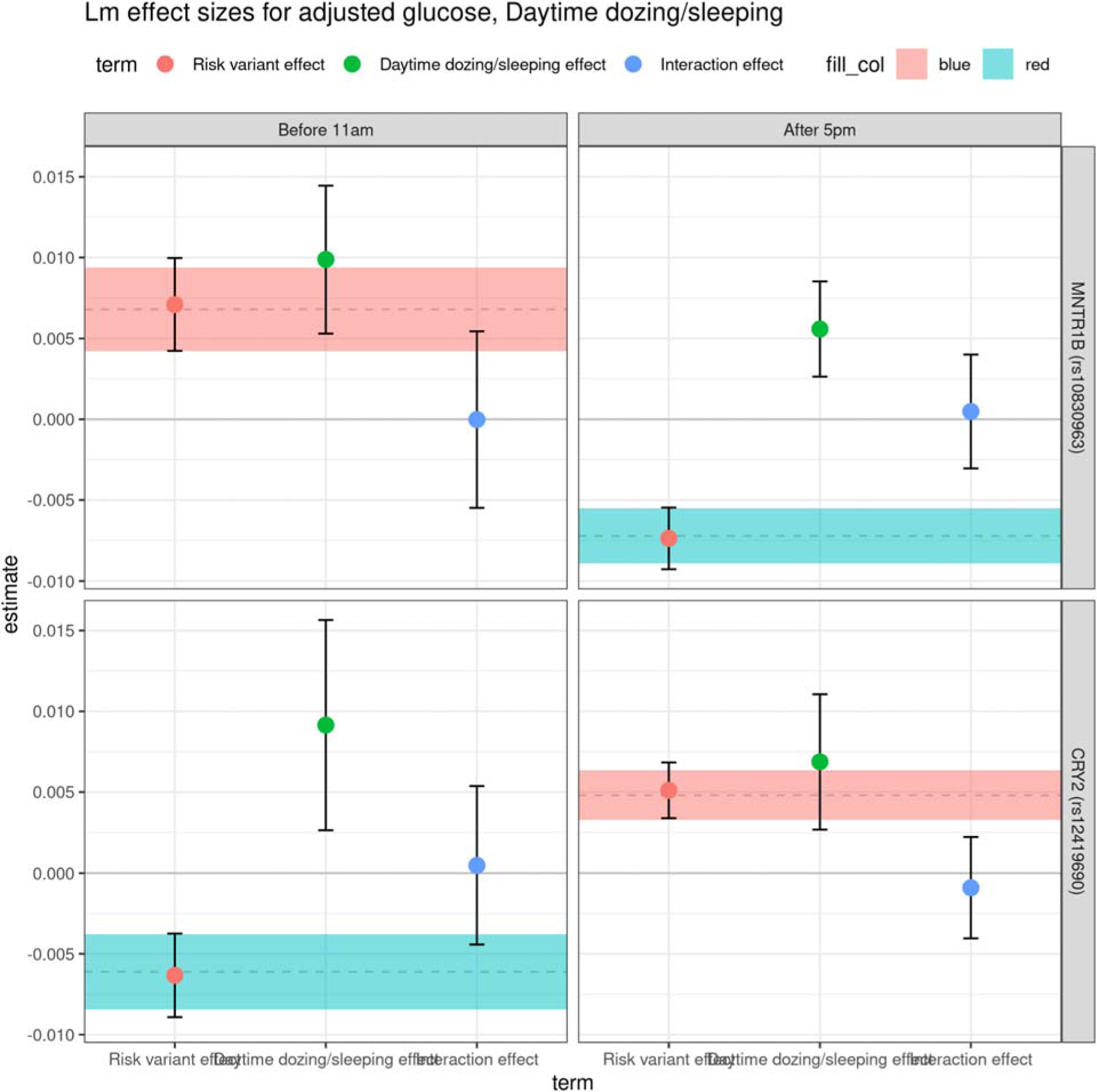
Effect of dozing, risk genotype and interaction between insomnia and genotype on glucose levels. We computed the association of dozing during the day (green), risk variant (red) and interaction effect (blue) on glucose levels. This shows an effect of dozing on glucose both in the morning and evening but no interaction between the genotype and dozing effect. Dashed lines and shaded area in background represent risk variant effect size and 95% confidence intervals over all samples.

**Supplementary Figure 16.**
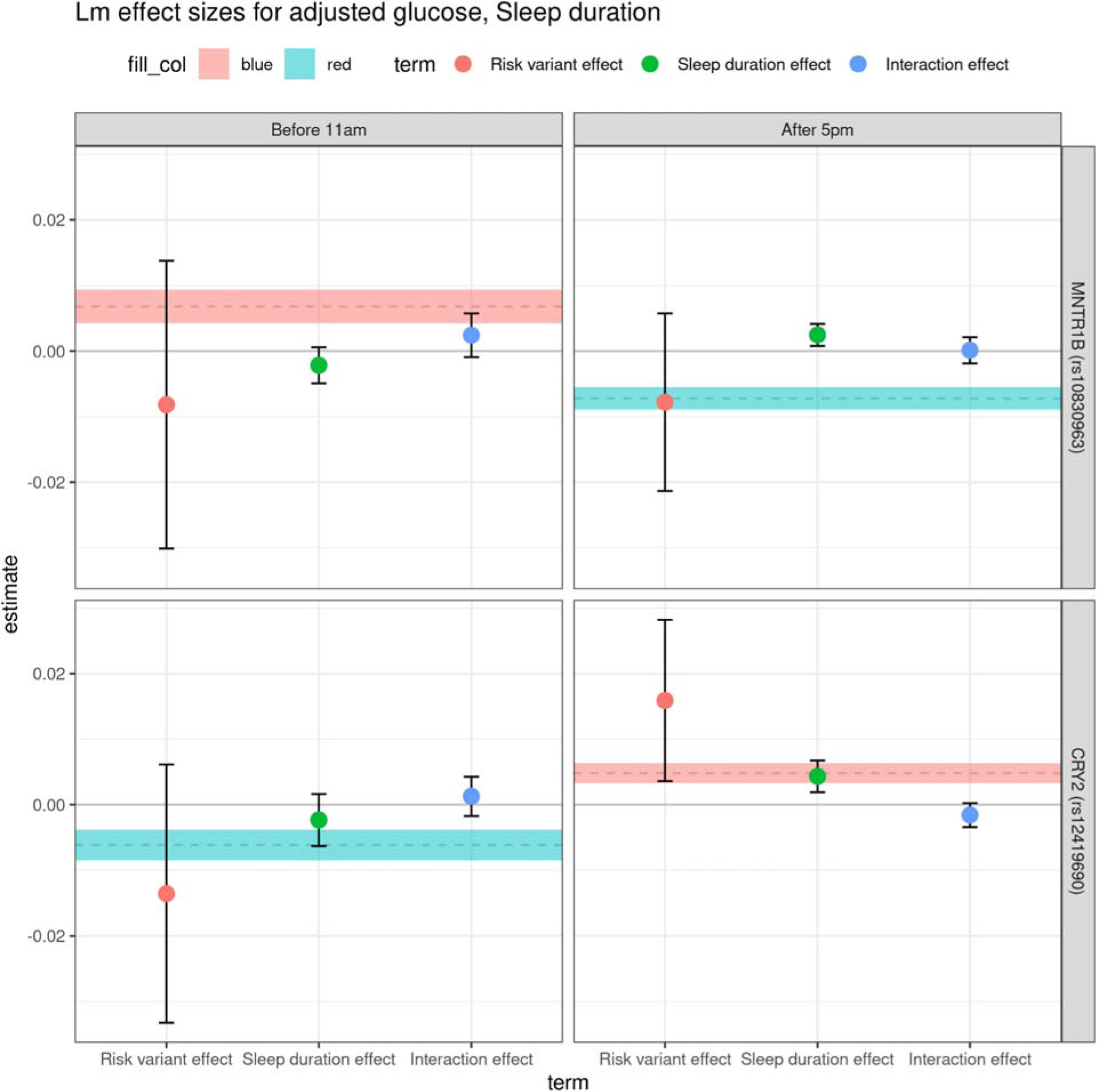
Effect of sleep duration, risk genotype and interaction between insomnia and genotype on glucose levels. We computed the association of sleep duration (green), risk variant (red) and interaction effect (blue) on glucose levels. Dashed lines and shaded area in background represent risk variant effect size and 95% confidence intervals over all samples.

**Supplementary Figure 17.**
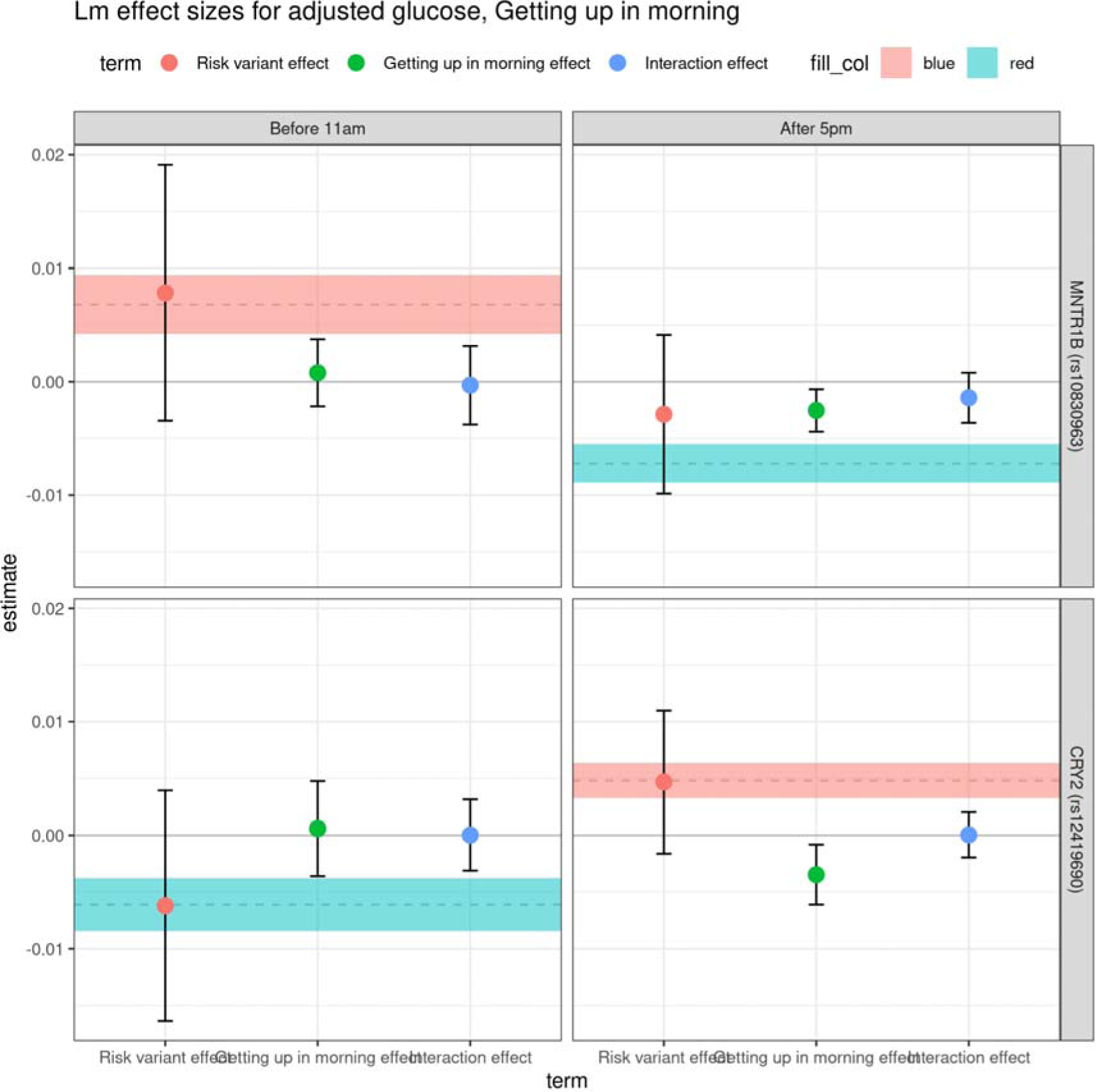
Effect of ease of awakening, risk genotype and interaction between insomnia and genotype on glucose levels. We computed the association of ease of awakening (green), risk variant (red) and interaction effect (blue) on glucose levels. Dashed lines and shaded area in background represent risk variant effect size and 95% confidence intervals over all samples.

## Acknowledgements

This work has been supported by NHGRI RM1HG010461 (NSA), The Finnish Foundation for Cardiovascular Research (HMO), and Instrumentarium Science Foundation (HMO).

This research has been conducted using the UK Biobank Resource under Application Number 22627 (University of Helsinki), 24983 (Stanford University) and 52374 (Fred Hutchinson Cancer Center). The UK Biobank study is funded by the following organizations: Medical Research Council, Wellcome Trust, Department of Health, Scottish Government, the Welsh Assembly Government, British Heart Foundation, Cancer Research UK and the Northwest Regional Development Agency.

The work of the Estonian Genome Center, University of Tartu was funded by the European Union through Horizon 2020 research and innovation program under grants no. 810645, 894987, 101137201 and 101137154, through the European Regional Development Fund project MOBEC008 and Estonian Research Council Grant PRG1291.

## Estonian Biobank Researchers

Andres Metspalu, Lili Milani, Reedik Mägi, Mari Nelis, Georgi Hudjashov,Tõnu Esko

### Ethics statement Estonian Biobank

The activities of the EstBB are regulated by the Human Genes Research Act, which was adopted in 2000 specifically for the operations of EstBB. Individual level data analysis in EstBB was carried out under ethical approval 1.1-12/624 from the Estonian Committee on Bioethics and Human Research (Estonian Ministry of Social Affairs), using data according to release application T19 6-7/GI/16001 from the Estonian Biobank.

### Ethics statement UK Biobank

The UK Biobank has received approval as a Research Tissue Bank from the North West Multi-centre Research Ethics Committee (MREC) under MREC permits 11/NW/0382 (2011–2016), 16/NW/0274 (2016–2021) and 21/NW/0157 (2021–2026). Researchers with approved applications are covered by these permits and are not required to seek additional approval, except in specific cases (see section B7 of the UK Biobank Access Procedures document: https://www.ukbiobank.ac.uk/media/omtl1ie4/access-procedures-2011-1.pdf). All participants of the UK Biobank study provided consent, at the baseline visit, for their personal data and biological samples to be collected and stored for research purposes. Participants are given the option to withdraw their consent at any time; any samples that have withdrawn their consent at the time of analysis were excluded from this study. A print version of the electronic consent form is stored as UK Biobank Resource 100252.

## REFERENCES

1 Laothamatas, I., Rasmussen, E. S., Green, C. B. & Takahashi, J. S. Metabolic and chemical architecture of the mammalian circadian clock. Cell Chem Biol 30, 1033–1052 (2023). 10.1016/j.chembiol.2023.08.014

2 Koronowski, K. B. & Sassone-Corsi, P. Communicating clocks shape circadian homeostasis. Science 371 (2021). 10.1126/science.abd0951

3 Ruben, M. D. et al. A database of tissue-specific rhythmically expressed human genes has potential applications in circadian medicine. Sci Transl Med 10 (2018). 10.1126/scitranslmed.aat8806

4 Scheer, F. A. & Czeisler, C. A. Melatonin, sleep, and circadian rhythms. Sleep Med Rev 9, 5–9 (2005). 10.1016/j.smrv.2004.11.004

5 Ahmad, S. B. et al. Melatonin and Health: Insights of Melatonin Action, Biological Functions, and Associated Disorders. Cell Mol Neurobiol 43, 2437–2458 (2023). 10.1007/s10571-023-01324-w

6 Scheer, F. A. et al. Repeated melatonin supplementation improves sleep in hypertensive patients treated with beta-blockers: a randomized controlled trial. Sleep 35, 1395–1402 (2012). 10.5665/sleep.2122

7 Bass, J. & Takahashi, J. S. Circadian integration of metabolism and energetics. Science 330, 1349–1354 (2010). 10.1126/science.1195027

8 Hujoel, M. L. A. et al. Protein-altering variants at copy number-variable regions influence diverse human phenotypes. Nat Genet 56, 569–578 (2024). 10.1038/s41588-024-01684-z

9 Henson, J. et al. Waking Up to the Importance of Sleep in Type 2 Diabetes Management: A Narrative Review. Diabetes Care 47, 331–343 (2024). 10.2337/dci23-0037

10 Schmid, S. M., Hallschmid, M. & Schultes, B. The metabolic burden of sleep loss. Lancet Diabetes Endocrinol 3, 52–62 (2015). 10.1016/S2213-8587(14)70012-9

11 Cappuccio, F. P. et al. Meta-analysis of short sleep duration and obesity in children and adults. Sleep 31, 619–626 (2008). 10.1093/sleep/31.5.619

12 Kalsbeek, A., la Fleur, S. & Fliers, E. Circadian control of glucose metabolism. Mol Metab 3, 372–383 (2014). 10.1016/j.molmet.2014.03.002

13 Shapiro, E. T. et al. Oscillations in insulin secretion during constant glucose infusion in normal man: relationship to changes in plasma glucose. J Clin Endocrinol Metab 67, 307–314 (1988). 10.1210/jcem-67-2-307

14 Sabatti, C. et al. Genome-wide association analysis of metabolic traits in a birth cohort from a founder population. Nat Genet 41, 35–46 (2009). 10.1038/ng.271

15 Lyssenko, V. et al. Common variant in MTNR1B associated with increased risk of type 2 diabetes and impaired early insulin secretion. Nat Genet 41, 82–88 (2009). 10.1038/ng.288

16 Prokopenko, I. et al. Variants in MTNR1B influence fasting glucose levels. Nat Genet 41, 77–81 (2009). 10.1038/ng.290

17 Dupuis, J. et al. New genetic loci implicated in fasting glucose homeostasis and their impact on type 2 diabetes risk. Nat Genet 42, 105–116 (2010). 10.1038/ng.520

18 Lagou, V. et al. GWAS of random glucose in 476,326 individuals provide insights into diabetes pathophysiology, complications and treatment stratification. Nat Genet 55, 1448–1461 (2023). 10.1038/s41588-023-01462-3

19 Elliott, A. et al. Distinct and shared genetic architectures of gestational diabetes mellitus and type 2 diabetes. Nat Genet 56, 377–382 (2024). 10.1038/s41588-023-01607-4

20 Mahajan, A. et al. Multi-ancestry genetic study of type 2 diabetes highlights the power of diverse populations for discovery and translation. Nat Genet 54, 560–572 (2022). 10.1038/s41588-022-01058-3

21 Jones, S. E. et al. Genome-wide association analyses of chronotype in 697,828 individuals provides insights into circadian rhythms. Nat Commun 10, 343 (2019). 10.1038/s41467-018-08259-7

22 Watanabe, K. et al. Genome-wide meta-analysis of insomnia prioritizes genes associated with metabolic and psychiatric pathways. Nat Genet 54, 1125–1132 (2022). 10.1038/s41588-022-01124-w

23 Lane, J. M. et al. Biological and clinical insights from genetics of insomnia symptoms. Nat Genet 51, 387–393 (2019). 10.1038/s41588-019-0361-7

24 Jansen, P. R. et al. Genome-wide analysis of insomnia in 1,331,010 individuals identifies new risk loci and functional pathways. Nat Genet 51, 394–403 (2019). 10.1038/s41588-018-0333-3

25 Dashti, H. S. et al. Genome-wide association study identifies genetic loci for self-reported habitual sleep duration supported by accelerometer-derived estimates. Nat Commun 10, 1100 (2019). 10.1038/s41467-019-08917-4

26 Sinnott-Armstrong, N. et al. Genetics of 35 blood and urine biomarkers in the UK Biobank. Nat Genet 53, 185–194 (2021). 10.1038/s41588-020-00757-z

27 Vetter, C. et al. Night Shift Work, Genetic Risk, and Type 2 Diabetes in the UK Biobank. Diabetes Care 41, 762–769 (2018). 10.2337/dc17-1933

28 Vetter, C. et al. Association Between Rotating Night Shift Work and Risk of Coronary Heart Disease Among Women. JAMA 315, 1726–1734 (2016). 10.1001/jama.2016.4454

29 Tuomi, T. et al. Increased Melatonin Signaling Is a Risk Factor for Type 2 Diabetes. Cell Metab 23, 1067–1077 (2016). 10.1016/j.cmet.2016.04.009

30 Lane, J. M. et al. Genome-wide association analyses of sleep disturbance traits identify new loci and highlight shared genetics with neuropsychiatric and metabolic traits. Nat Genet 49, 274–281 (2017). 10.1038/ng.3749

31 Donga, E. & Romijn, J. A. Sleep characteristics and insulin sensitivity in humans. Handb Clin Neurol 124, 107–114 (2014). 10.1016/B978-0-444-59602-4.00007-1

32 Sudlow, C. et al. UK biobank: an open access resource for identifying the causes of a wide range of complex diseases of middle and old age. PLoS Med 12, e1001779 (2015). 10.1371/journal.pmed.1001779

33 Ahmadi, M. N., Huang, B. H., Inan-Eroglu, E., Hamer, M. & Stamatakis, E. Lifestyle risk factors and infectious disease mortality, including COVID-19, among middle aged and older adults: Evidence from a community-based cohort study in the United Kingdom. Brain Behav Immun 96, 18–27 (2021). 10.1016/j.bbi.2021.04.022

34 Batty, G. D., Gale, C. R., Kivimaki, M., Deary, I. J. & Bell, S. Comparison of risk factor associations in UK Biobank against representative, general population based studies with conventional response rates: prospective cohort study and individual participant meta-analysis. BMJ 368, m131 (2020). 10.1136/bmj.m131

35. Leitsalu, L., et al. Cohort Profile: Estonian Biobank of the Estonian Genome Center, University of Tartu. Int J Epidemiol 44, 1137-1147 (2015). 10.1093/ije/dyt268

36 Allen, N. E. et al. Approaches to minimising the epidemiological impact of sources of systematic and random variation that may affect biochemistry assay data in UK Biobank. Wellcome Open Res 5, 222 (2020). 10.12688/wellcomeopenres.16171.2

37 Wain, L. V. et al. Novel insights into the genetics of smoking behaviour, lung function, and chronic obstructive pulmonary disease (UK BiLEVE): a genetic association study in UK Biobank. Lancet Respir Med 3, 769–781 (2015). 10.1016/S2213-2600(15)00283-0

38 McCarthy, S. et al. A reference panel of 64,976 haplotypes for genotype imputation. Nat Genet 48, 1279–1283 (2016). 10.1038/ng.3643

39 Huang, J. et al. Improved imputation of low-frequency and rare variants using the UK10K haplotype reference panel. Nat Commun 6, 8111 (2015). 10.1038/ncomms9111

40 Bycroft, C. et al. The UK Biobank resource with deep phenotyping and genomic data. Nature 562, 203–209 (2018). 10.1038/s41586-018-0579-z

41 Mitt, M. et al. Improved imputation accuracy of rare and low-frequency variants using population-specific high-coverage WGS-based imputation reference panel. Eur J Hum Genet 25, 869–876 (2017). 10.1038/ejhg.2017.51

42 Browning, B. L., Tian, X., Zhou, Y. & Browning, S. R. Fast two-stage phasing of large-scale sequence data. Am J Hum Genet 108, 1880–1890 (2021). 10.1016/j.ajhg.2021.08.005

43 Manichaikul, A. et al. Robust relationship inference in genome-wide association studies. Bioinformatics 26, 2867–2873 (2010). 10.1093/bioinformatics/btq559

44 Mbatchou, J. et al. Computationally efficient whole-genome regression for quantitative and binary traits. Nat Genet 53, 1097–1103 (2021). 10.1038/s41588-021-00870-7

